# Keto-gluconeogenic metabolic axis mirrors renal homeostasis and post-injury response in spatial transcriptomics

**DOI:** 10.64898/2026.09.24.753087

**Authors:** Paul Gueguen, Judith Vouillamoz, Safia Hadjadj, Kateryna Spahic, Roland H. Wenger, Stellor Nlandu Khodo

**Affiliations:** Functional Genomics Center Zurich, University of Zurich and ETH Zurich, Zurich, Switzerland; Department of Physiology, Faculty of Medicine and Sciences, University of Zurich, Zurich, Switzerland; National Institute of Health and Medical Research (INSERM), UMR-S1155, Tenon Hospital, Faculty of Medicine, Sorbonne University, Paris, France

**Author notes:** Address correspondence to: Dr. Stellor Nlandu Khodo, PhD INSERM UMR S1155, CoRaKid, Bâtiment Recherche Tenon Hospital, 4, rue de la Chine 75020 Paris, FRANCE.

**Keywords:** kidney, acute kidney injury, chronic kidney injury, post-injury renal regeneration, renal metabolism, short chain fatty acid metabolism, ketone body metabolism, spatial transcriptomics

## Abstract

Metabolic disturbance is a key feature in acute kidney injury (AKI) and chronic kidney disease (CKD). The kidney highly depends on fatty acid metabolism; how renal cells rewire metabolism upon AKI and CKD transition remained unclear. Here, we combine spatial transcriptomics with biochemical analyses to characterize metabolic changes and intercellular interactions during AKI to CKD transition. Using aristolochic acid model of CKD, we spatio-temporally correlated changes in structure and function with metabolic profile during AKI to CKD transition. Surprisingly, keto-gluconeogenic metabolic axis was highly enriched in healthy kidney and paralleled injury phase transitions. Intercellular interaction analysis associated a subtype of macrophage with the drift to chronicity. Online scRNAseq datasets analysis confirmed altered keto-gluconeogenic metabolic pathways in ischemia-reperfusion injured mouse kidneys and AKI and CKD human kidney biopsies compared to healthy controls. This study identifies a metabolic axis transcriptionally mirroring renal homeostasis and injury responses, and tubulo-interstitial interactions that can be targeted to mitigate AKI to CKD transition.

---

The kidney is a highly metabolic organ endowed with restricted post-injury regenerative capacity and metabolic disturbance is a major factor in post-injury regeneration failure^1–4^. Renal macroscopic anatomy comprises the highly vascularized and oxygen dependent cortex and less vascularized and relatively hypoxic medulla. Each renal structural and functional unit, the nephron, includes a filter, the glomerulus, and the tubular system composed of the proximal tubule (PT S1, S2 and S3), the loop of Henle (LOH), the distal tubule (DT) connected to the collecting duct (CD) by the connecting tubule (CNT). Each segment tunes metabolism according to its function, tubulo-vascular topology and microenvironment^5–7^. Renal parenchyma remains quiescent under normal physiology; however, upon an acute kidney injury (AKI)-induced abrupt loss of function, epithelial cells undergo cell-autonomous mechanisms, rewire their metabolism and communicate with interstitial cells (stromal/immune cells) to restore near to normal architecture and function^8–14^. Most compelling studies reported the pivotal role of injured PT in both post-injury repair process and the drift toward chronic kidney disease (CKD)^15, 16^. Depending on the injury frequency and severity, survived PT cells dedifferentiate, express progenitor markers such as Sox9, FoxM1, proliferate and re-differentiate to restore renal structure and function^17, 18^. PT cells dwell in the cortex where oxygen availability meets their high oxidative metabolism to produce required energy for active transport activities^6, 19^. We recently reported that thick ascending limb (TAL), a crucial part of LOH and PT cells are the most vulnerable renal segments partly due to their high dependency to mitochondrial and fatty acid metabolism to produce energy, and defective epithelial fatty acid metabolism contributes in kidney fibrosis development^20–22^. Moreover, in contrast to other renal epithelial cells, PT cells have the particularity to not only use but also generate glucose through gluconeogenesis^23–29^. Though the kidney reportedly utilizes long chain and medium chain fatty acid and amino acid to produce energy^30^, increasing number of studies suggested the beneficial role of gut microbiota derived short chain fatty acids (SCFA) in renal response to injury^31–34^. These SCFAs notably butyrate and propionate are not only fatty acid metabolic intermediates, but they may also play a role in epigenetic gene regulation and injury repair^35–37^. Among epithelial pathogenic metabolic changes, promotion of anaerobic glycolysis is well described upon AKI and CKD^38^. However, how metabolic changes mirror post-AKI recovery and the transition to CKD is not well understood. Using histological and spatial transcriptomic approaches, we designed a model of post-AKI recovery and transition to CKD in mice through repetitive injections of aristolochic acid (AA) and transcriptionally analyzed intercellular interactions and how post-injury metabolic rewiring correlates with peak acute (3 weeks), transition (5 weeks) and chronic injury (8 weeks) phases. This study depicted the limited regenerative capacity of the kidney post-acute injury and CKD transition using histology and spatial transcriptomics and suggests a key role of cortical keto-gluconeogenic metabolism in renal homeostasis.

## Results

### Post-injury incomplete regeneration and restored renal function failed to impede CKD

The kidney restores renal structure and function post-AKI and CKD transition suggests collapsed regenerative capacity. To investigate post-injury renal regenerative capacity, mice were injured with aristolochic acid (AA) and analyzed at different time points corresponding to peak acute injury (AKI), transition (recovery) and chronic (CKD) phases. HE and Qupath Eosin channel histological analysis confirmed peak acute injury at 3 weeks time point (3W) characterized by increased tubular injury (dilation, flattening, and atrophy), interstitial cell infiltrate and decreased tubular mass (Figure 1a, b). After strong injury at the peak acute phase, as expected, kidneys showed tissue restoration reflected by decreased tubular injury, interstitial infiltrate and increased tubular mass as compared to the peak acute phase; however, it was incomplete as compared to uninjured kidney tissue (Figure 1a, b). Despite recovery attempt observed at 5 weeks time point (5W), kidneys became more injured with significant cellular infiltrate and reduced tubular cortico-medullary mass, suggesting a collapse of repair process at 8 weeks time point (8W) (Figure 1a, b). Analysis of cortical collagen accumulation on Sirius red stained kidney slices confirmed the transition to chronicity at 8W (Figure 1c, d). Collagen accumulation was expectedly not statistically different at the peak acute (3W), transition (5W) and uninjured phase (Figure 1c, d). Renal function was strikingly decreased in the peak acute injury phase and restored close to normal at the recovery/transition phase (5W), even though renal architecture did not reach normality compared to uninjured kidneys (Figure 1a, b, e, f). Despite apparent tubulo-interstitial fibrosis, renal function (BUN and ACR) was lower at 8W as compared to peak acute phase (3W), implying a non-linear temporal correlation between fibrogenesis and function impairment in chronic phase (8W) (Figure 1c-f). Taken together, this result confirmed restricted regenerative capacity of the kidney post-AKI which, for non-clearly elucidated reasons, collapses leading to fibrosis upon chronic injury. Moreover, this result suggests potential function/structure correlation inadequacy in CKD classification post-AKI.

**Figure 1:**
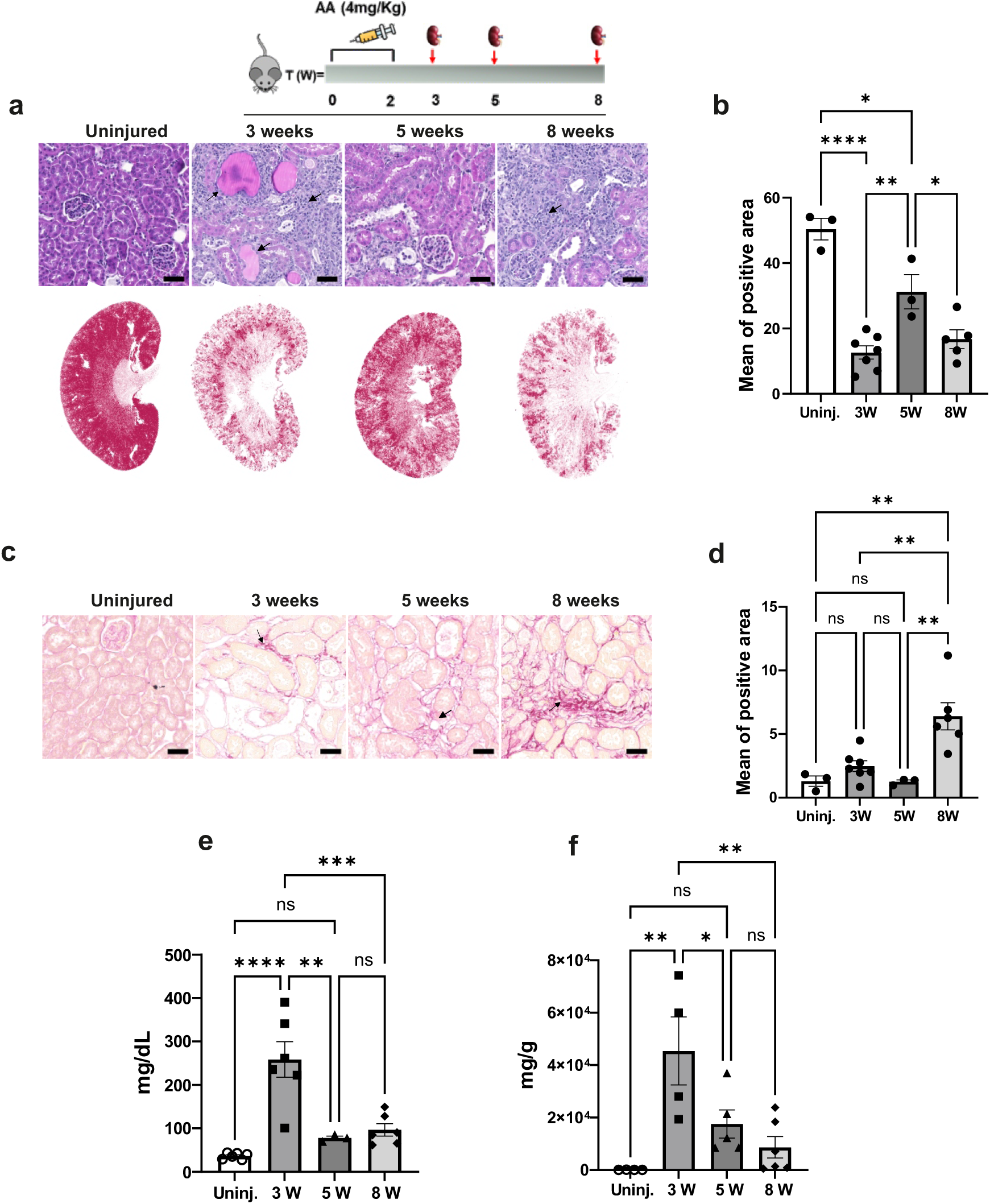
Insufficient post-acute injury regeneration leads to fibrogenesis. (a) Representative H&E (upper panel) and Qupath Eosin channel (lower panel) images of kidneys from uninjured, 3 weeks, 5 weeks and 8 weeks AA injured mice. Arrows indicate injured tubules (dilatation, flattening, atrophy and luminal cast deposition) and interstitial cell infiltrate. (b) Quantification of Qupath Eosin channel red area; n=3 (uninjured), n=7 (3W), n= 3 (5W), and n= 5 (8W) mice. (c) Representative Sirius red staining images of kidneys from uninjured, 3 weeks, 5 weeks and 8 weeks AA injured mice showing collagen accumulation in red. Arrows indicate fibrotic areas. (d) Quantification of Sirius red positive area; n=3 (uninjured), n=7 (3W), n= 3 (5W), and n= 6 (8W) mice, p= (test). (e) Plasma blood urea nitrogen (BUN) levels measured at 0 (uninjured), 3 weeks, 5 weeks and 8 weeks AA injur y time points 6 weeks; n=6 (uninjured), n= 6 (3W), n= 3 (5w) and 6 (8w) AA injured mice, p=0. (test). (f) Urine albumin-creatinine ratio (ACR) levels measured at 0 (uninjured), 3 weeks, 5 weeks and 8 weeks AA injur y time points 6 weeks; n=4 (uninjured), n= 4 (3W), n= 4 (5W) and 6 (8W) AA injured mice, p=0. (test). All scale bars (a and c) represent 100mm; dots represent the number of animals per group (b, d, e and f). Data are presented as mean values ±SEM. Statistical significance was determined by one way ANOVA followed by multiple comparisons test with p<0.05 considered statistically significant unless otherwise stated * represents p-value<0.05; ** represents p-value <0.01; **** represents p-value <0.0001; ns= nonsignificant.

### Spatial transcriptomics confirmed incomplete regeneration despite restored renal function

To transcriptionally characterize acute to chronic injury transition as defined in histology and estimate cell type proportion per spot, we performed spatial transcriptomics (Visium) on uninjured and injured kidneys and applied robust cell type decomposition (RCTD)^39^ algorithm using curated reference dataset for deconvolution (GSE197266). Analysis of mRNA expression of injury (Vcam1), myofibroblast (Acta2) and fibrosis (Col1a1) markers confirmed peak acute, transition and chronic injury phases at 3W, 5W and 8W injury time points respectively (Figure 2a). After cell deconvolution, mRNA expression of renal segment specific markers resolved on uninjured kidney slice displayed normal spatial localization and confirmed high proportion of PT cell positive spots in the cortex and outer medulla (Slc5a2+, Slc22a8+, Slc13a3+ and Slc22a7+) while LOH cells were enriched in the inner medulla (Umod+), suggesting a correct cell type assignment and deconvolution (Figure 2b and Extended Data Figure 1). Deconvolved cell type proportions resolved on HE stained renal slides revealed impaired renal cortico-medullary organization upon renal injury as compared to uninjured kidneys and confirmed post-acute injury regeneration attempt in the transition phase (5W) (Figure 2c, upper panel). The most vulnerable PT S3 cells were correctly enriched in deep cortex/outer medulla and were drastically decreased at the peak acute injury (3W), only the proportion of PT S3 dwelling in the outer medulla survived at the peak acute phase in accordance with our previous studies^10^. Consistent with our histological result, the proportion of PT cells increased in transition phase (5W) compared to the peak acute injury phase and decreased in chronic injury phase (8W) (Figure 2c and Extended Data Figure 2, lower panel). Cell type proportion analysis and multiple comparison using conditional false discovery rate confirmed previous studies where PT cells contribution to total mRNA in whole kidney transcriptomic data was greater than 50% [Clark, 2019] and indicated cellular remodeling upon injury (Figure 2d). PT (S1, S2 and S3) segment proportion is decreased in injury phases. While the S1 and S3 segments are drastically decreased, notably S3, the S2 segment proportion tends to increase in injury phase, suggesting a possible intra-segment remodeling or differentiation (loss of specific markers) notably S1/S3 to S2 segment (Figure 2d and Extended Data Figure 2). As observed in Figure 2c, S3 PT proportion was strikingly decreased in the acute phase, partially recovered in transition phase and decreased again in the chronic phase compared to uninjured. S1 and S1/S2 PT cell proportion was also decreased in the acute phase, partially recovered in the incomplete regeneration phase and decreased again in the chronic phase compared to uninjured. The proportion of injured PT cells was enhanced in the peak acute injury phase.

**Figure 2:**
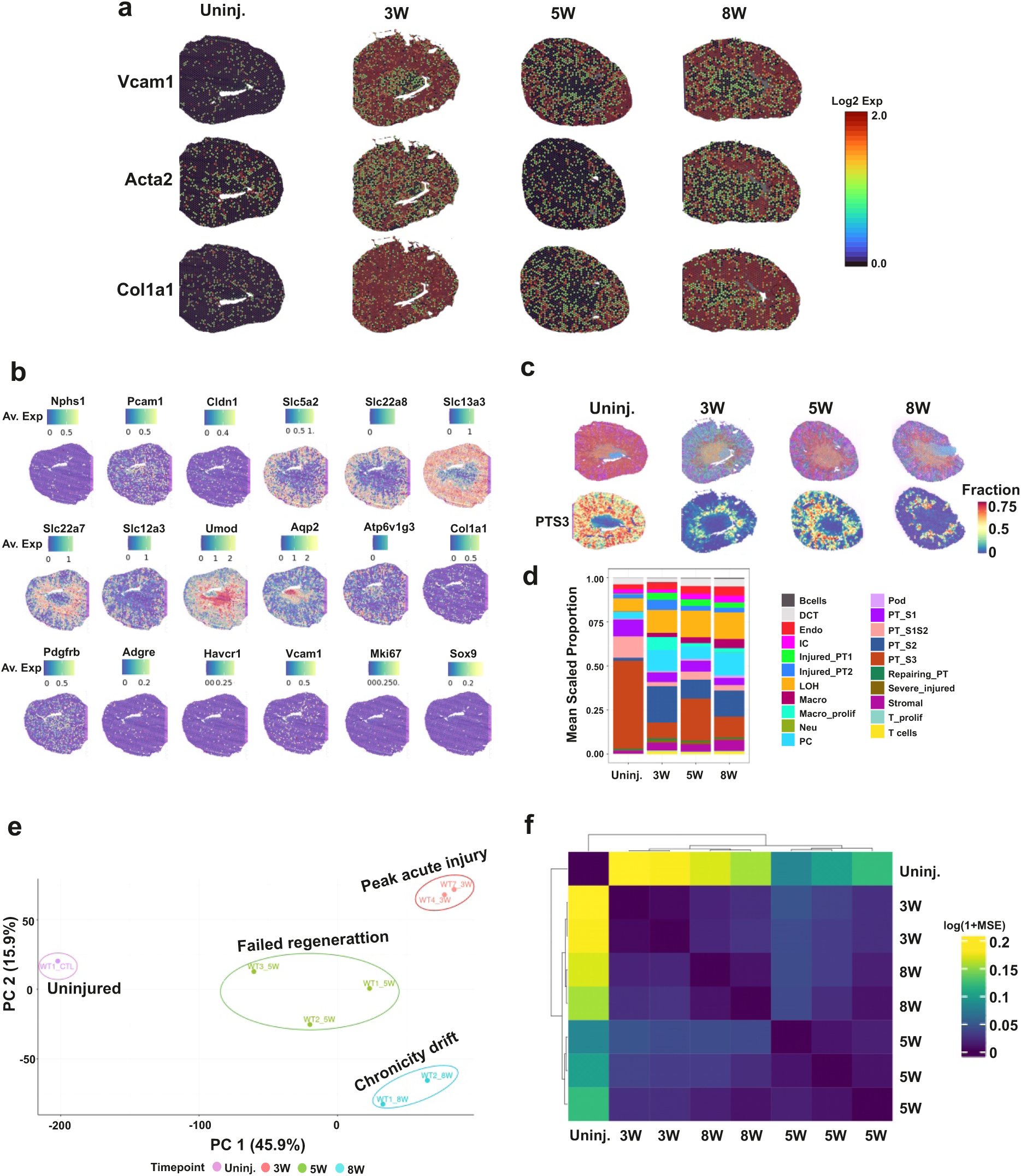
Spatial transcriptomics characterization of acute to chronic injury transition. (a) Spatial transcriptomics Loupe browser kidney images showing proximal tubule injury (Vcam1), myofibroblast (Acta2) and stromal (Col1a1) markers at baseline and throughout the injury phases. (b) Representative spatial featplots of selected renal segment specific markers resolved on uninjured kidney slices. (c) Post-deconvolution representative images of cell type pie chart resolved on HE stained tissue slides (upper panel). Representative images of post-deconvolution PT S3 cells resolved on uninjured, 3W, 5W and 8W injured kidney slices (lower panel). (d) Post-deconvolution cell type proportion plot per condition. DCT, distal convoluted tubule cells; Endo, endothelial cells; IC, collecting duct intercalated cells; Injured_PT1, type 1 injured proximal tubule cells; Injured_PT2, type 2 injured proximal tubule cells, LOH, loop of Henle cells; Macro, macrophages; Macro_prolif, proliferating macrophages; Neu, neutrophiles; T_prolif, proliferating T cells; PC, collecting duct principal cells; Pod, podocytes; PT, proximal tubule cells; S1/S2/S3, segments 1/2/3; Severe injured, severely injured PT; Stromal, stromal cells. (e) Pseudo-bulk Principal component analysis plot of uninjured and injured kidneys. (f) Heatmap representing the mean squared error between samples. n= 1 (uninjured), n= 2 (3W), n= 3 (5W) and n= 2 (8W).

While injured_PT2 (Havcr+, Krt8+, Rpl4+, Vcam^low^ and Ccl5^low^,) proportion was approximately divided by 2, proportion of injured_PT1 (Havcr+, Lcn2+, Fn1+, Vcam^high^ and Ccl5^high^) remained the same in the transition phase, suggesting their potential role in the progression to chronic phase. Apart from PT cells, PC and LOH proportion increased in injury phase. PC proportion partially decreased in the transition phase compared to peak acute injury phase and increased in the chronic phase back to the peak acute level, whereas LOH cells proportion gradually increased through the course of injury, suggesting a dynamic tubular remodeling upon injury (Figure 2d). Stromal cell proportion was increased upon injury and higher in the chronic injury phase, implying their potential role in fibrogenesis. Almost absent in uninjured kidney, injury triggered immune cell (Macrophages and T cells) proportion. Macrophages proliferation and polarization (M1/M2) are associated with post-injury recovery and the drift to chronicity^40–42^. Macrophage proportion significantly varies throughout injury phases, and we identified two subsets of macrophages, a proliferative subpopulation (Adgre1+ and Mki67+) and non-proliferating macrophages (Adgre1+, Cd86+, Adgre1+ and Arg1+). The proportions of proliferating macrophages (Macro_prolif) and other macrophages (other_Macro) are significantly increased in peak acute injury phase compared to uninjured, and Macro_prolif were higher compared to Macro (Figure 2d). Interestingly, the proportion of Macro_prolif was significantly decreased in the incomplete regeneration transition phase while other_Macro proportion was slightly increased as compared to peak acute injury phase. Moreover, Macro_prolif remained low whereas other_Macro were increased in chronic phase. Taken together, this result suggests a potential role of Macro_prolif. in peak acute injury which is switched off in incomplete regeneration transition phase. T cells were induced in injury without big difference throughout the course of injury (Figure 2d). Pseudo-bulk principal component analysis (PCA) of spatial transcriptomic data distinctly clustered samples and corroborated the three injury states identified in histology (Figure 1). Uninjured samples clustered at the left (low PC1) whereas injured samples tend toward the right part of the diagram (high PC1). 3W injured (peak acute phase) samples were (high PC1, low PC2) whereas 8W samples (chronic phase) were (high PC1, high PC2). 5W samples in the transition phase displayed an intermediate state between uninjured and 3W/8W samples, being high PC1 compared to uninjured samples but low PC1 compared to 5W and 8W samples (Figure 2e). Moreover, mean squared error (MSE) distance metrics confirmed similarities (lower distance) between uninjured and 5W injured kidneys; whereas 3W and 8W samples were less similar (high distance) compared to uninjured (Figure 2f). This transcriptional result corroborated our histological findings and clearly suggests an incomplete regeneration post-acute injury which for some unknown reasons drifts to chronic phase.

### Downregulation of keto-gluconeogenic metabolism in injured kidneys

Cell metabolism dictates the pace of the injury response, and metabolic disturbances are associated with acute and chronic injury. To transcriptionally determine metabolic pathways associated with the three predefined kidney injury phases, we performed pathway analyses across conditions and cell types. KEGG pathway analysis of differentially expressed genes indicated peroxisome, carbon, fatty acid and amino acid metabolisms as the most downregulated pathways in injury phases whereas inflammatory processes were upregulated in injured phase compared to uninjured phase (Extended Data Figure 3). Though these metabolic pathways were downregulated upon injury, UCell score metabolic pathway enrichment analysis revealed partial recovery in the transition phase (5W) compared to peak acute injury phase (3W) which collapsed in chronic injury phase (8W) (Extended Data Figure 4). Surprisingly, among the most affected metabolic pathways paralleling the pattern of injury phases (downregulated in peak acute injury phase, incompletely restored in transition phase and downregulated in chronic phase) we found previously reported pathways such as fatty acid β-oxidation, gluconeogenesis, amino acid degradation, tricarboxylic acid (TCA) cycle^4^, but also short chain fatty acid (SCFA) metabolism (propionate and butyrate metabolism), ketogenesis, branched amino-acid metabolism (valine, leucine and isoleucine degradation), glyoxylate and dicarboxylate metabolism and peroxisome function (Figure 3a, b and Extended Data Figure 4). To exclude the effect of injury model, we analyzed an online scRNAseq dataset (GSE139107) of ischemia reperfusion injury (ctl. 4h, 12h, 2 days, 14 days and 6 weeks)^16^. Keto-gluconeogenesis axis is significantly impaired throughout injury course (Extended Data Figure 5). Interestingly, these metabolic pathways were also enriched in PT cells, particularly the S3 segment, as compared to the rest of the tubular system, and decreased in injured PT cells (Extended Data Figures 4 and 6). Consistent with PT cells enrichment, spatial resolution of enriched metabolic pathways on H&E-stained kidneys revealed cortical localization (Figure 3c). Fatty acid metabolism impairment was also reflected by increased lipid deposition in renal tissue during the peak acute injury phase (not statistically significant) and chronic phase (Figure 3d).

**Figure 3:**
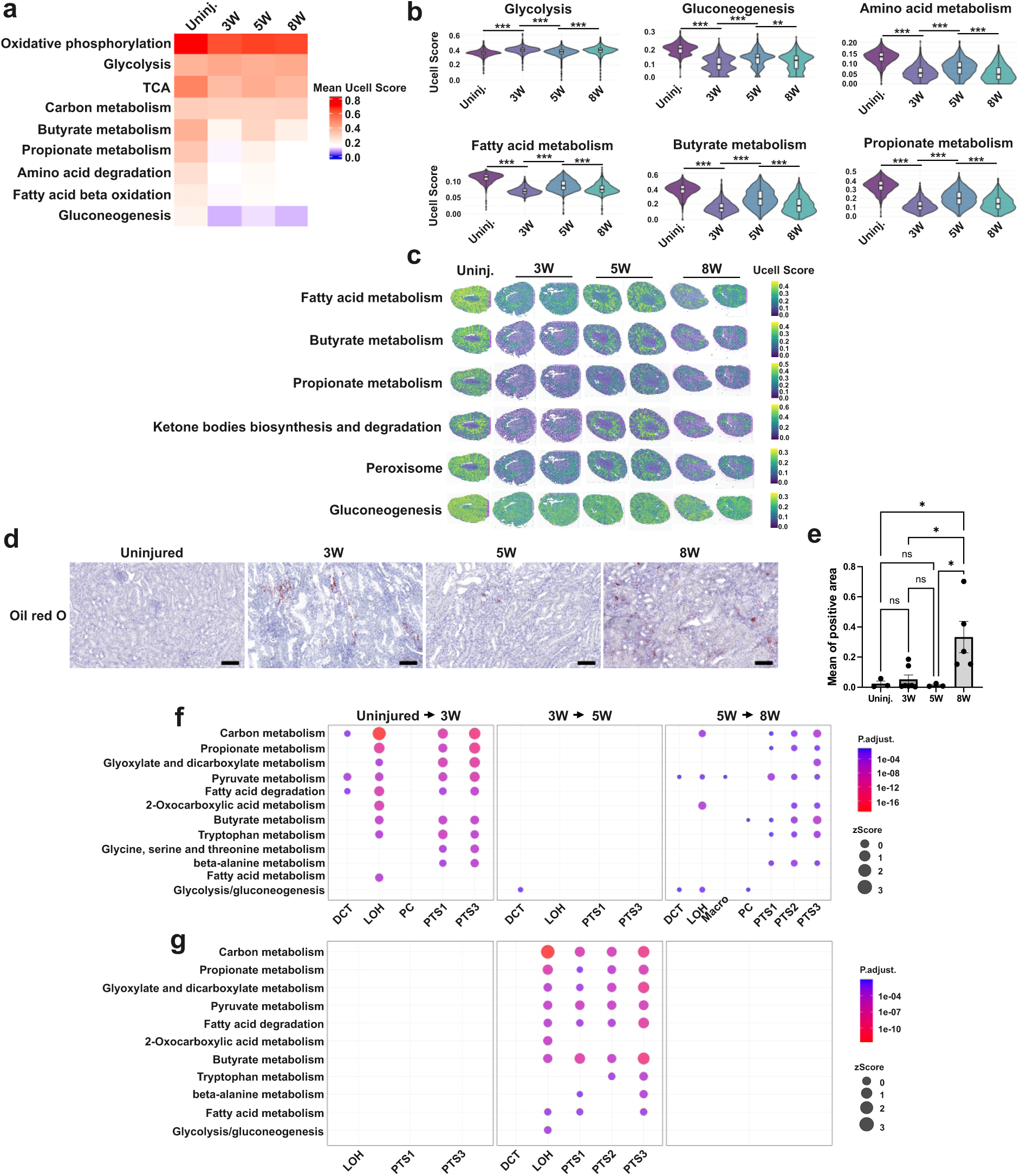
Prominent role of keto-gluconeogenic metabolic pathways in renal homeostasis and post-injury recovery. (a) UCell score heatmap showing enrichment of the main oxidative metabolic pathways between uninjured and injured conditions. (b) UCell score violin plots showing the main cortical metabolic pathways between uninjured and injured kidneys. Statistical significance of pathway differences was determined by Kruskal-Walli’s test across injury phases (p < 2.10^-16^). (c) Spatial representation of keto-gluconeogenic pathway enrichment resolved on H&E-stained kidney slices. (d) Representative images of oil red O-stained uninjured and injured kidneys. (e) Quantification of oil red O-stained kidneys. n = 3 (uninjured); n = 7 (3W); n = 3 (5W) and n = 5 (8W). Statistical significance was determined by one-way ANOVA followed by multiple comparisons test with p < 0.05 considered statistically significant. *, p < 0.05; **, p < 0.01; ****, p <0.0001; ns = non-significant. Scale bars are 100 μm. (f) Comparative analysis of high z-score metabolic pathways throughout the course of injury, showing upregulated pathways from uninjured to 8W injury phase. (g) Comparative analysis of high z-score metabolic pathways throughout the course of injury showing downregulated pathways from uninjured to 8W injury phase. DCT, distal convoluted tubule cells; LOH, loop of Henle cells; PC, collecting duct principal cells; PT, proximal tubule cells; S1/S2/S3, segments 1/2/3.

Comparative pathway analysis throughout renal epithelia during injury phases confirmed impairment of keto-gluconeogenic metabolic pathways and identified LOH and PT as the main metabolically adapting nephron segments in the transition phase (Figure 3f, g). LOH cells showed higher enrichment in carbon metabolism, including carbohydrate metabolism. LOH sub-clustering analysis identified 4 clusters (0, 1, 2 and 3). Though enriched in TAL markers, notably in cluster 0 (Slc12a1+ and Kcnj1+), analysis of specific markers suggests a mixture of TAL, distal tubules, collecting duct, urothelial cells, some outer medullary PTs and immune cells, suggesting that the metabolic signature of the LOH involved more than one cell type (Extended Data Figure 7). Surprisingly, while the kidney reportedly prefers long and medium-chain fatty acids in energy metabolism^30^, SCFA metabolism in PT and LOH, notably propionate and butyrate metabolism, was strongly enriched and upregulated in uninjured phase compared to 3W time point (Figure 3f) whereas SCFA metabolism was downregulated in 3W compared to 5W time points (Figure 3g). Comparison of these pathways between 5W and 8W confirmed their incomplete restoration in the transition phase particularly in the PT and LOH segments (Figure 3f, g).

Taken together, these results imply a pivotal role of keto-gluconeogenic metabolism in renal tissue homeostasis. Incomplete metabolic restoration during the post-AKI transition phase is insufficient to sustain normality and impedes CKD transition.

### Short-chain fatty acids promote proximal tubule mitochondrial function

To refine the list of metabolic pathways associated with incomplete regeneration during the transition phase, we analyzed differentially expressed genes and identified 463 signature genes in the transition phase (Figure 4a and Supplementary table S1). Gene ontology analysis using EnrichR identified mitochondria, intracellular organelles and peroxisomes as the most affected cellular components, suggesting a dysfunction of the mitochondria-peroxisomal metabolic axis, which may be required for adaptive fatty acid and amino acid metabolism (Figure 4b). Among these keto-gluconeogenic pathways re-induced in the transition phase, the “Synthesis and degradation of ketone bodies” term has the highest odds ratio followed by the “Valine, leucine and isoleucine degradation”, “Butyrate metabolism”, “Propionate metabolism”, “Glyoxylate and dicarboxylate metabolism” and “Peroxisome” terms (Figure 4c).

**Figure 4:**
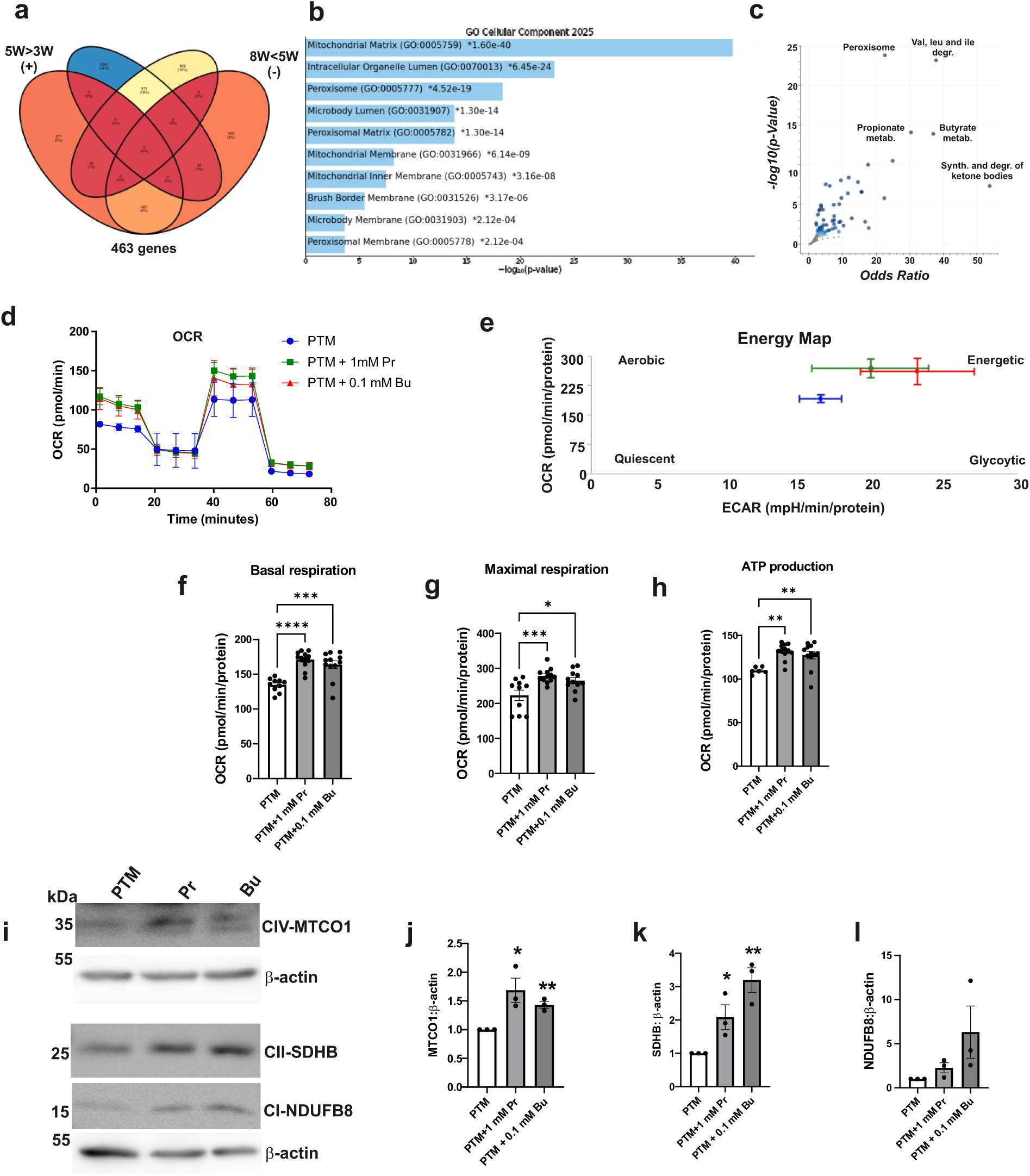
Propionate and butyrate promote proximal tubule mitochondrial function. (a) 4 ways Venn diagram showing common differentially expressed genes across conditions. (b) Bar chart showing top 10 enriched metabolic terms in KEGG database based on common 463 genes characterizing the transition phase as compared to peak acute and chronic injury phases. Colored bars correspond to terms with significant p-values (<0.05). An asterisk (*) next to p-value indicates the term also has a significant adjusted p-value (<0.05). (c) Volcano plot of KEGG metabolic terms depicting odds ratio and -log(p-value) of enriched metabolic terms from the 463 input query gene set. (d) Seahorse Mito-stress test showing oxygen consumption rate variation at baseline and maximal respiration in PT media (PTM) or treated with 1mM propionate (Pr) or 0.1mM butyrate (Bu). (e) Seahorse energy map plot showing the metabolic status (aerobic, quiescent, glycolytic and energetic) of untreated and propionate and butyrate treated PT cells. (f) Quantification of basal oxygen consumption rate between untreated and propionate and butyrate treated PT cells. (g) Quantification of maximal oxygen consumption rate between untreated and propionate and butyrate treated PT cells. (h) Quantification of ATP- linked OCR production between untreated and propionate and butyrate treated PT cells (n=3 independent biological replicates). Data are presented as mean values ±SEM. Statistical significance was determined by one way ANOVA followed by multiple comparisons test with p<0.05 considered statistically significant unless otherwise stated * represents p-value<0.05; ** represents p-value <0.01; **** represents p-value <0.0001; ns= nonsignificant. (i) Representative images of immunoblots showing expression of mitochondrial complex (I, II and IV) subunits and β-actin in untreated and propionate and butyrate treated PT cells. (j, k, l) Quantification of MTCO1, SDHB and NDUFB8 immunoblots. Data are presented as mean values ±SEM. Statistical significance was determined by unpaired Student’s *t* test (two groups) with p<0.05 considered statistically significant. * represents *p*<0.05; ** represents *p*<0.01; **** represents *p*<0.0001.

Butyrate and propionate are fatty acid metabolic intermediates involved in energy production, reported to act as epigenetic regulatory elements linking diet, metabolism and gene expression^35^. Therefore, we treated PT cells with propionate and butyrate, and found increased mRNA levels of enzymes involved in gluconeogenesis and ketogenesis (Extended Data Figure 8). Moreover, PT cells treated with propionate and butyrate increase basal and maximal oxygen consumption rates (OCR) as well as ATP production, making PT cells more energetic and aerobic compared to untreated cells (Figure 4d-h). Surprisingly, propionate and butyrate treatment increased mitochondrial complex II and IV subunits.

In conclusion, these results associate incomplete regeneration attempts during the transition phase with re-induction of keto-gluconeogenic metabolic pathways and demonstrated crucial roles of butyrate and propionate in proximal tubule energy production, mitochondrial function and promotion of ketogenic and gluconeogenetic metabolism.

### Differential epithelial-interstitial cell Interactions throughout injury phases

Upon injury, parenchymal cells undergo autonomous changes, interact with neighboring cells, and activate interstitial cells to restore tissue architecture and function. Immune and stromal cells actively participate in the repair process, and both lymphocytes and macrophages are increased early after injury^43^. To determine cell-cell interactions, notably epithelial cells interacting with macrophage and stromal subpopulations throughout injury phases, we performed cell-cell communication analyses using the cell chat database of ligand-receptor pairs^44^. Cellchat-infered ligand-receptor connectivity increased after injury compared to the uninjured state (Figure 5a). The number of total inferred interactions was lower during the peak acute phase as compared to the transition and chronic phases; the number of interactions were quite similar in transition and chronic phases (Figure 5b, c).

**Figure 5:**
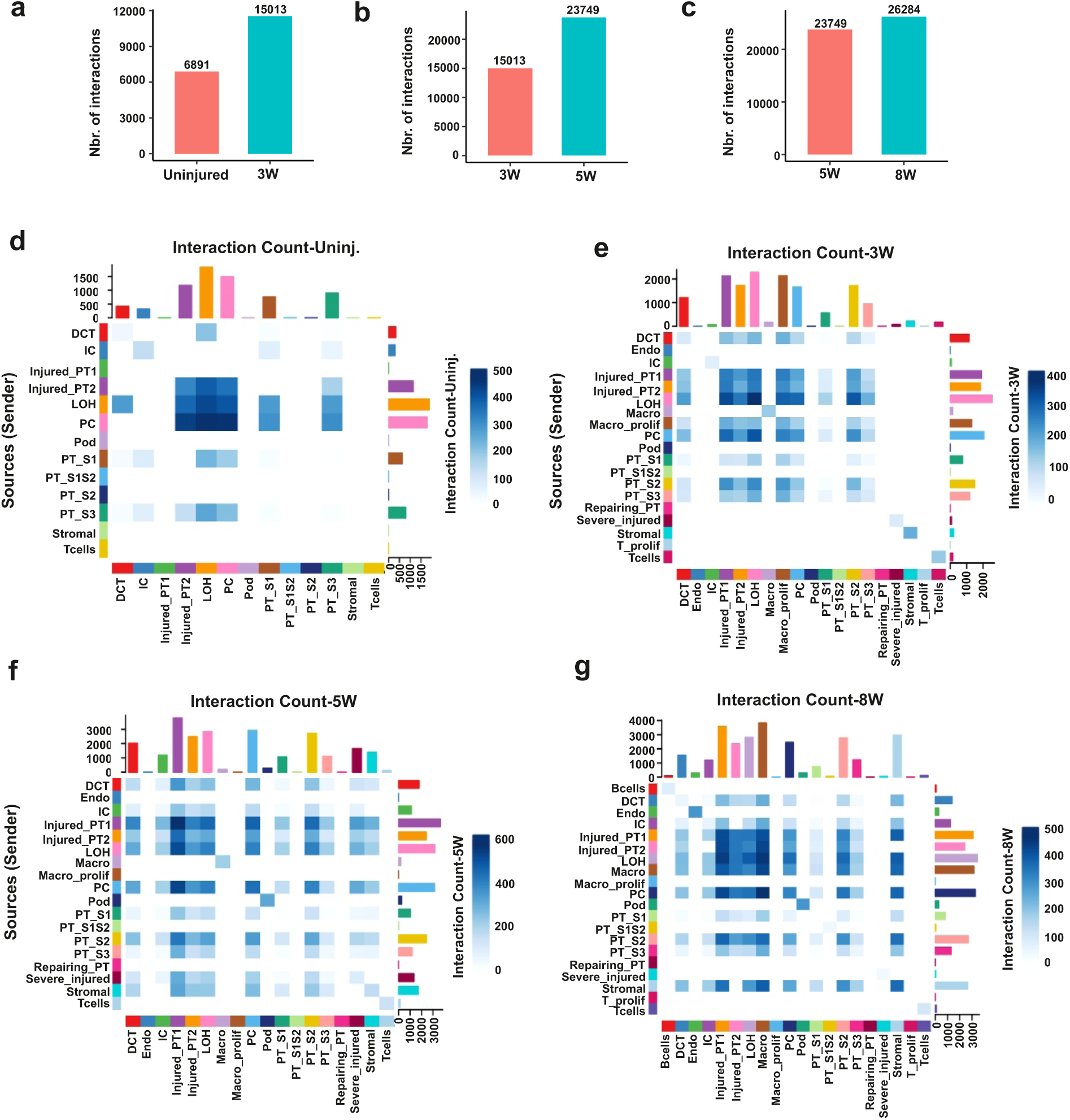
Differential intercellular interactions across conditions. (a-c) Histograms showing comparisons of the number of total interactions between uninjured and peak acute injury phases (a), peak acute injury and transition phases (b), and transition and chronic injury phases (c). (d-g) Cell chat heat maps of cell type interactions showing cell type interactions in uninjured (d), peak acute injury (e), transition (f) and chronic injury phases (g). DCT, distal convoluted tubule cells; Endo, endothelial cells; IC, collecting duct intercalated cells; Injured_PT1, type 1 injured proximal tubule cells; Injured_PT2, type 2 injured proximal tubule cells, LOH, loop of Henle cells; Macro, macrophages; Macro_prolif, proliferating macrophages; Neu, neutrophiles; T_prolif, proliferating T cells; PC, collecting duct principal cells; Pod, podocytes; PT, proximal tubule cells; S1/S2/S3, segments 1/2/3; Severe injured, severely injured PT; Stromal, stromal cells.

In uninjured kidneys, LOH, collecting duct/CD (PC and IC) and injured_PT2 were the most interacting cell types throughout renal epithelia (Figure 5d). During peak acute injury, in addition to LOH, PC and injured_PT2, PT cells (S1, S2 and S3) and injured-PT1 showed autocrine and paracrine interactions with renal epithelia, including with LOH, PC, injured PT and Macro_prolif. Renal epithelial cells, notably uninjured and injured PT, LOH and PC, interacted with Macro_prolif but not with other_Macro which showed autocrine interaction despite their increased proportion compared to uninjured kidney (Figure 2c and Figure 5e). Given their abundance, PT cells (uninjured and injured) had more interactions with Macro-prolif, suggesting an important role of PT cells during post-injury activation of macrophages. Macro_prolif mainly interacted with PT and LOH segments whose metabolisms were significantly impacted through the injury phases, notably in the peak acute injury phase (Figure 3f, g and Figure 5e). Moreover, no inferred intercellular interactions were mediated by stromal cells during the peak acute injury phase, despite strong induction of myofibroblast and fibrotic marker transcript levels (Figure 2a and Figure 5e). Surprisingly, these epithelial-Macro_prolif interactions disappeared in the incomplete regeneration transition phase.

Neither proliferating nor non-proliferating macrophages interacted with epithelial cells at this stage (Figure 2c and Figure 5f). In contrast to the peak acute phase, epithelial-stromal interactions were promoted in the transition phase, though fibrotic marker transcripts tended to decrease (Figure 2a and Figure 5f).

Overall, the transition phase was characterized by enhanced epithelial cell interactions, absence of immune cell interaction (macrophages and T cells) with other cells, as well as absence of stromal-epithelial cell interactions. During the chronic phase, the number of other_Macro-mediated interactions with epithelial and stromal cells increased, while the number of Macro_prolif that did not mediate any interactions was reduced (Figure 2c and Figure 5g). Stromal cell interactions increased, notably with injured PT1, PTS2, LOH, PC and Macro (Figure 5g). To understand the role of Macro_prolif and other_Macro, UCell score M1/M2 signature enrichment analysis was performed and revealed enrichment of the M1 signature in Macro_prolif compared to other_Macro, whereas other_Macro showed an enriched M2 signature compared to Macro_prolif, suggesting distinct roles of these two populations through the injury phases (Extended Data Figure 9).

Taken together, these results suggest differential interactions between the most metabolically impacted LOH and PT segments with stromal cells and macrophages during injury phase transition.

### Decreased keto-gluconeogenic metabolism in human AKI and CKD

Impaired metabolism is common during AKI and CKD progression^45^. Cross-species validation was performed using harmonized single-cell atlas that encompasses healthy controls and diseased human kidney biopsies. We investigated keto-gluconeogenic pathway enrichment across disease states, including normal, acute kidney failure (AKF), and chronic kidney disease (CKD)^46^. UCell score analyses revealed impaired keto-gluconeogenic metabolism in AKF and CKD compared to healthy kidney biopsies. The keto-gluconeogenic pathways identified in mice were reduced in AKI and CKD compared to healthy controls (Figure 6). Moreover, these pathways were not only impaired in renal epithelial cells, but also in interstitial cells (Extended Data Figure 10), implying for a complex tubulo-interstitial metabolic rewiring during AKF and CKD transition.

**Figure 6:**
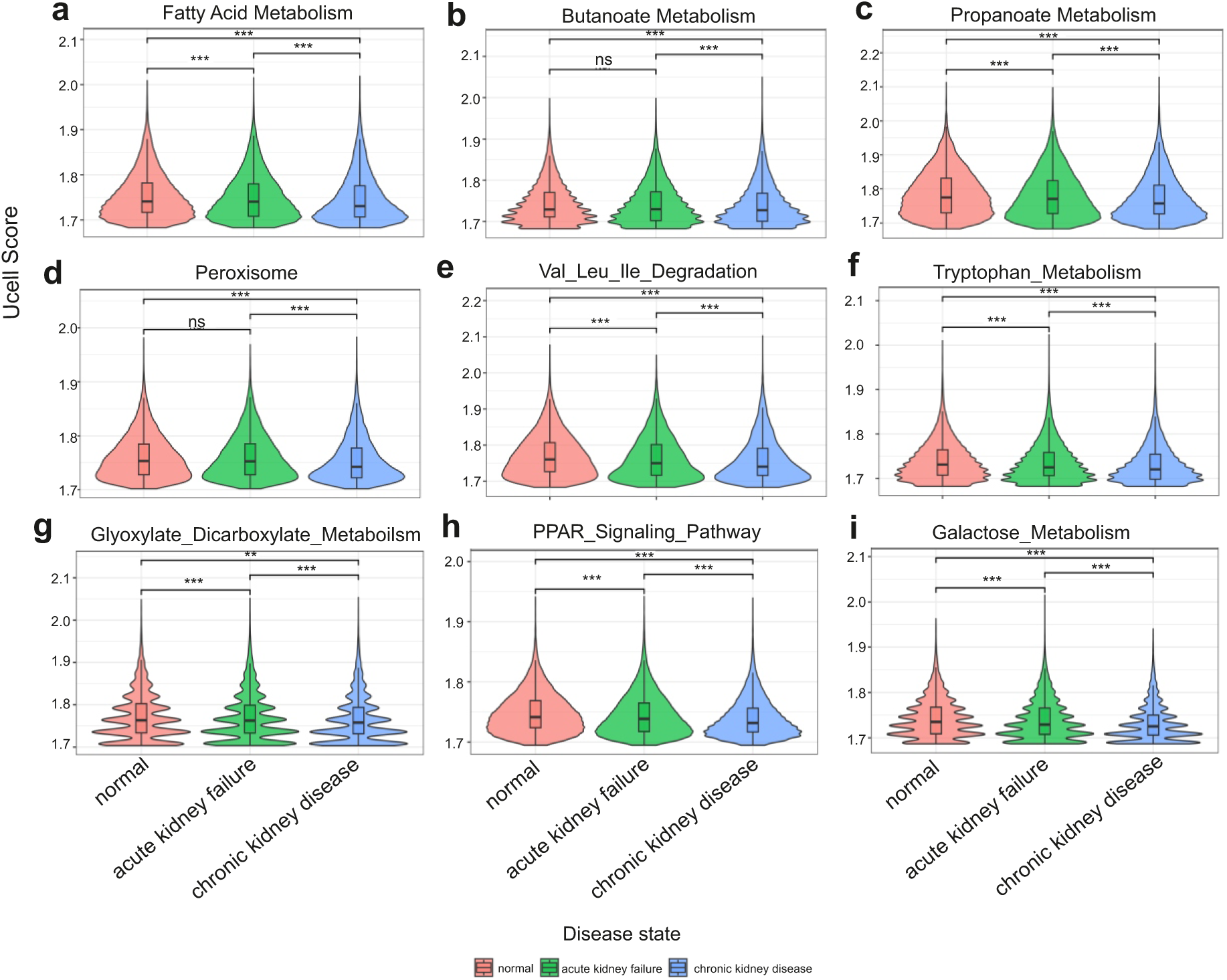
Decreased keto-gluconeogenic metabolic pathways in AKI and CKD. UCell violin plots showing fatty acid metabolism (a), butanoate metabolism (b), propanoate metabolism (c), peroxisome (d), valine,leucine and isoleucine degradation (e), tryptophan metabolism (f), glyoxylate and dicarboxylate metabolism (g), PPAR signaling pathway (h) and galactose metabolism (i) pathways enrichments in healthy controls, AKF and CKD kidney biopsies from integrated single-cell atlas of healthy and injured human kidney biopsies, enabling comparisons across disease states (GSE121862). Comparisons were statistically assessed using Wilcoxon rank-sum tests (*, p < 0.05; **, p < 0.01; ***, p < 0.001).

## Discussion

High metabolic rates and dependency on oxidative metabolism are key features of the kidney. However, the special renal tubulo-vascular topography imposes a cortico-medullary oxygen consumption gradient and metabolic substrate preference difference throughout the tubular system. While highly perfused cells in the renal cortex are dependent on oxidative phosphorylation, the relatively hypoxic cells dwelling in the inner medulla mostly rely on anaerobic glycolysis. Several studies linked cellular metabolism rewiring with post-injury survival and the drift to chronic injury^3, 21, 47^. The PT is the most sensitive renal segment to injury, partly due to its high metabolic rate, oxygen dependency and exposure to toxins. We recently demonstrated how the dependency on oxidative metabolism, notably on fatty acid β-oxidation, sensitizes PT and TAL cells to AKI^20^. Given the pivotal role of PT cells in post-AKI regeneration and CKD transition, we investigated how this high metabolic and sensitive hub to injury rewires metabolism in post-injury recovery and during the transition of AKI to CKD, using spatial transcriptomics and biochemical approaches. After structural and functional assessments of renal integrity, we defined time points corresponding to peak acute, incomplete regeneration/transition and chronic phases upon aristolochic acid nephrotoxicity.

At peak acute injury, we observed a catastrophic renal injury, reflected by impaired structure (tubular injury, interstitial infiltrate and reduced parenchyma), which was confirmed by an abrupt decline in renal function. During the transition window, renal injury was partially resolved as reflected by decreased injury compared to the peak acute injury phase and restoration of renal function. Despite partial structural and functional renal recovery, kidneys have drifted to fibrogenesis at the chronic phase, reflected by collagen accumulation and sustained decrease of renal function. Spatial transcriptomic profiling of metabolic pathways identified carbon metabolism and peroxisome as the most dysregulated metabolic pathways in acute and chronic injury phases. Interestingly, keto-gluconeogenic metabolic pathways were the most decreased in the peak acute injury phase and incompletely restored in the transition phase. Among fatty acid metabolic pathways, SCFAs metabolism, notably propionate and butyrate metabolism in PT and LOH cells, correlated the most with post-injury recovery during the transition phase, though incompletely restored and collapsed in the chronic injury phase. Moreover, treatment of conditionally immortalized PT cells with propionate and butyrate increased the OCR implying for homeostatic effect of SCFAs in renal metabolism.

Both histology and spatial transcriptomics confirmed post-AKI regenerative capacity of the kidney and the drift to CKD. Though mRNA data do not always reflect protein data, our study confirmed the adequacy of assessing renal injury on the mRNA level. Given the low resolution of Visium, we used the RCTD method to deconvolve renal cell types which has the advantage over Seurat label transfer by providing multi-cell modeling, quantitative proportions, corrections for systematic technical differences between single-cell RNA-seq references and Visium spatial transcriptomics, while Seurat’s anchor-based integration assumes similar feature spaces. The deduced cell type proportions, notably for PT cells, were similar to prior reports^48^. Cell type proportion analyses between uninjured and injured phases confirmed a model of partial post-injury recovery and transition to CKD. KEGG pathways analyses using differentially expressed genes indicated decreased traditional substrate metabolism and increased inflammatory pathways throughout injury phases compared to uninjured kidneys. Whereas most studies reported fatty acid metabolism as the most affected metabolic pathways in PT cells^21^, we identified mainly cortical keto-gluconeogenic metabolism as the most affected metabolic pathway during acute to chronic injury transition.

The kidney is known to utilize long chain and medium chain fatty acids as well as amino acids to produce energy. While recent studies reported the beneficial effect of gut microbiota-derived SCFAs in renal response to injury^37^, the high scores of SCFA (propionate and butyrate metabolism) and branched chain amino-acids (BCAAs) in uninjured kidneys suggest a potential cell autonomous mechanism in the kidney promoting this exotic metabolic axis. Consistently, “peroxisome” metabolic axis, which plays a central role in fatty acid breakdown and biosynthesis^49^, was highly enriched in uninjured kidney and strongly affected upon injury. Whether peroxisomes of cortical cells, notably of PT cells, act as checkpoints controlling the length of the fatty acid chains to be metabolized by mitochondria, remains to be investigated. These SCFA metabolic pathways tend to be restored during the incomplete regenerative transition phase, supporting their association with renal homeostasis and recovery. The transient decrease in preferred substrate metabolism may represent a survival mechanism to reduce oxygen consumption; however, depending on injury severity, this survival mechanism might promote maladaptive processes, including G2/M cycle arrest, leading to chronic injury and fibrosis^50^. The decrease of keto-gluconeogenic metabolic axis during the chronic phase reflects impairment of renal homeostatic and regenerative capacity, likely due to persistent maladaptive tubulo-interstitial interactions and enhanced anaerobic metabolic pathways that deviate recovered keto-gluconeogenic metabolic axis and promotes CKD.

Comparative metabolic analyses throughout the renal epithelia identified PT and LOH cells as the most metabolically impacted cell types. Though we expected PT cells to be metabolically affected by kidney injury, it is difficult to conclude about the LOH based on our sub-clustering data which revealed a mixture of different cell types. However, LOH was enriched with TAL markers which have been reported to be an active hub of fatty acid oxidation.

Several studies reported beneficial roles of gut-microbiota derived SCFAs upon AKI and CKD; other studies reported the role of SCFAs in gene regulation by acting on chromatin accessibility^35, 51^. Propionate and butyrate treatment augmented PT cells’ oxidative metabolic capacity and increased the expression of mitochondrial complex II and IV subunits, suggesting that these SCFAs may not only act as metabolites but also play a role in mitochondria functional homeostasis. Propionate and butyrate metabolisms promote ketone metabolism and gluconeogenesis by providing metabolic intermediates or regulating expression of specific enzymes involved this metabolic axis^52, 53^. Unlike propionate which directly enters gluconeogenic pathway, butyrate necessitates an activation step into butyryl-CoA prior to being used for glucose synthesis. Moreover, butyrate acts indirectly by promoting expression of gluconeogenic genes^54, 55^. Butyrate stimulates the production of ketone bodies through induction of key enzymes involved in ketogenesis^56, 57^.

Cell-cell interaction analyses identified differential intercellular interaction through the injury phases. Given their abundance, PT cells were at the center of intercellular interactions, however, LOH and PC cells were also mediating important intercellular interactions with injured and uninjured PT cells, macrophages and stromal cells.

Inflammation is a common feature of AKI, which depending on injury severity and frequency, may participate in post-injury recovery or promote progression to CKD. Proliferating macrophages exclusively interacted with renal epithelial cells at peak acute injury while the remaining population of macrophages rather mediate epithelial and stromal interaction during the chronic phase. Proliferating macrophages, showing a M1 signature, might mediate inflammation and be switched off during the incomplete recovery transition phase, while other macrophages, showing a M2 signature, might participate in the repair process between the transition and chronic phases. Previous studies reported a double-edged sword effects of M2 macrophage-mediated repair and fibrogenesis through excessive production of ant-inflammatory and profibrotic factors such as TGF-β1^58^. Since proliferating macrophages are not pure M1 macrophages, they may mediate adaptive inflammatory mechanisms during the peak acute injury phase that lead to incomplete regeneration during the transition phase. Their unresponsiveness/inactivation later in the transition phase may promote the drift to CKD. One remaining question is whether macrophage-renal epithelial cell-cell interactions are influenced by the metabolic status of epithelial cells. Though out of scope of this study, future studies should investigate how parenchymal cell metabolism influences post-injury epithelial-interstitial cell interactions. Despite enhanced fibrotic marker mRNA expression, we did not observe fibrosis in histology at peak acute phase, suggesting that fibrotic marker mRNA induction likely reflects post-injury repair processes.

In the analyzed human databases, keto-gluconeogenic pathways are reduced in AKF and CKD renal biopsies. Unfortunately, signatures of these pathways cannot be investigated in post-AKI recovery state due to lack of data. Not only PT cells but also interstitial cells showed keto-gluconeogenic metabolic changes, suggesting a complex tubulo-interstitial metabolic rewiring upon AKF and CKD.

In conclusion, we confirmed restricted regenerative capacity of injured kidney and show that the keto-gluconeogenetic metabolic axis mirrors renal homeostasis, post-AKI regenerative capacity and the drift to CKD. The SCFAs propionate and butyrate appear to play an important role in this axis by modulating mitochondrial function and regulating the expression of genes involved in mitochondrial electron transport chain and keto-gluconeogenic axis. Tubulo-interstitial cell interactions vary through the injury phases, and missing interactions of a subpopulation of macrophages during the transition phase may be crucial for AKI to CKD transition. While most studies reported fatty acid metabolism as most affected in PT and TAL injury, our study provides a more precise metabolic axis that can be targeted to develop novel therapeutic strategies to mitigate AKI to CKD transition.

## Methods

### Animal models

All procedures were approved by the veterinary office of the canton Zurich, Switzerland (ZH123/19). Mice were tagged using ear notching in accordance with the Laboratory Animal Services Center (LASC) license 101, and the generated tissues were used for genotyping. After genotyping, animals were transferred to the experimental room, where they were allowed to acclimate for at least 7 days before starting experiments. During the acclimation period, mice were randomly assigned as controls or CKD and monitored to ensure water and food ad libitum accessibility every other day. Given the pathogenic strain of AKI to CKD models, the painkillers were used to minimize procedure-related pain. Pure FVB mice (Tgfbr2^fl/fl^)^59^ were injured (six repetitive intraperitoneal injections of 3 mg/kg aristolochic acid (AA; Sigma-Aldrich, location) every other day in two weeks) or not (uninjured) and kidneys were collected at 0 (uninjured), 3 (peak acute injury), 5 (transition phase) and 8 (chronic phase) weeks for histological, biochemical and spatial transcriptomics experiments. Mice were monitored every other day during 2 weeks of intraperitoneal AA injections, then every day in the acute phase (7 days after the last AA injection) and thereafter every other day until euthanasia. Mice were scored for signs of pain (hunched posture, poor grooming, reduced mobility and subsequent body weight loss) every day during the acute phase. According to the pain scoring criteria, mice were provided with wet food pellets and/or administered (ip injection) pre-warmed Ringer’s lactate/5% glucose solution and/or buprenorphine (0.1 mg/kg) diluted in 0.9% NaCl (1 ml of 0.3 mg/ml of buprenorphine in 5 ml of 0.9% NaCl, resulting in 2 μl/g body weight). Buprenorphine is an opioid and strong analgesic that we preferred in this study because of its long-lasting effect (6-8 h) and compared to other opioids (butorphanol for instance), it reportedly has minimal hemodynamic side effects which is a very important aspect in this study. Euthanasia was considered in case of failure of pain mitigating measures. If euthanasia is needed before the experimental endpoint, mice were killed by 70% CO_2_ exposure and only the kidneys were collected for further investigation. At the experimental endpoint, mice were anesthetized by inhalation of 5% isoflurane in oxygen using the VetFlo stand (Rothacher medical Gmbh).

After confirmation of complete anesthesia by checking the pedal withdrawal reflex and tail pinch three times, mice were killed by cervical dislocation followed by organ removal. The personal phone number of the study director and the experimenter, including their designed substitutes, were purposely put on the animal ID card to be contacted for emergency intervention to avoid animals suffering.

### AA Injury Models

We intraperitoneally (ip) injected Tgfbr2^fl/fl^ mice (N10 FVB background) with 3 mg/kg aristolochic acid (AA) (Sigma-Aldrich) a total of six times over 2 weeks (ref). The mice were sacrificed at 3, 5 and 8 weeks of AA injection protocol.

### Reagents and antibodies

Reagent and antibody’s information is listed in the supplementary table 3 and 4 respectively.

### Tissue staining and injury score

Kidneys were harvested, fixed in 10% formalin, embedded in paraffin or OCT, sectioned (machine, company, location) and stained with hematoxylin and eosin (H&E), Picrosirius red or oil red O-stained kidney sections were scanned using Zeiss Axio Scan (company, location). Images were quantified with ZEN software (Zeiss) and batch processing was performed using Image J (FIJI, location). For quantification, 10 high power fields (HPFs) were taken per sample and positive areas were determined in a blinded fashion using Image J. For Picrosirius red, oil red O, and eosin-stained sections 10 high power fields or whole images (eosin) of kidney cortices were taken per sample, and stained areas were quantified using Image J.

### Blood urea nitrogen

Prior euthanasia, whole blood was collected from mice, placed in heparinized tubes and centrifuged to collect plasma. Blood urea nitrogen (BUN) levels were determined by the Zurich Integrative Rodent Physiology (ZIRP) facility of the University of Zurich.

### Urine albumin creatine ratio

Urine was collected from mice after euthanasia by involuntary micturition reflex or collected directly from the bladder using insulin syringe, placed in 1.5 ml centrifuge tubes and albumin creatine ratio (ACR) was determined by the ZIRP facility.

### Spatial transcriptomics

Mouse kidneys were harvested from 1 uninjured, 2 AA-3 weeks injured, 3 AA-5 weeks injured and 2 AA-8 weeks injured mice. The 8 samples were embedded in OCT compound (ref.458; Tissue-Tek,, SAKURA, location) in a cryo-mold, snap frozen in liquid nitrogen using a container with methyl butane, and immediately stored at −80°C. Thin sections (10 μm) were cut using a cryostat (CM 3050S; LEICA, location), and immediately transferred on the capture area of the 10X Genomics (location) gene expression slides. Tissues were equilibrated for 1 minute at 37°C, formalin-fixed for 30 minutes at −20°C in pre-cooled methanol, and H&E stained. Images were captured with the Zeiss Axio Scan microscope. After image acquisition, tissues were permeabilized, and reverse transcription and library construction were performed according to the manufacturer’s instructions (10x Genomics Visium Spatial Gene Expression FFPE workflow). Libraries were sequenced on an Illumina NovaSeq X Plus instrument, generating 150 bp paired-end reads per library. Raw sequencing data were processed using Space Ranger (v3.0.1; 10x Genomics) to perform demultiplexing, alignment, barcode and UMI counting, and quality control. Reads were aligned to the mouse reference genome GRCm39 (Ensembl Release M31). Space Ranger outputs, including filtered spot-by-gene count matrices and summary quality metrics, were used for downstream analysis.

### Preprocessing, normalization, and clustering

All downstream analyses were performed in R (v4.5.0). Seurat (v5.3.1.1000) was used to import the Space Ranger matrices, manage spot-level metadata, and perform standard preprocessing. Data were normalized using LogNormalize. Highly variable features were identified with Seurat defaults, and data were scaled prior to principal component analysis (PCA). Low-dimensional embeddings were generated by computing PCA followed by uniform manifold approximation and projection (UMAP), utilizing the first 30 principal components (PCs 1-30) for neighbor graph construction and UMAP generation. Neighbor graphs were constructed with FindNeighbors using dims=1:30, and clusters were resolved with FindClusters at resolutions specified in the figure legends. Anatomical region annotations (for example, cortex and medulla) were incorporated into the metadata to enable region-stratified summaries where indicated. For aggregation and display purposes, principal cells (PC) and PT-PEC were merged into a unified “PC” class to simplify comparisons that focus on functional trends across nephron compartments.

### Cell type deconvolution of Visium spots

Spatial cell type deconvolution was performed using RCTD v2.0 (Cable et al., 2022). We used a mouse kidney single-cell RNA-seq atlas (GSE197266) as reference, comprising 28 cell types after filtering for minimum 50 cells per type and downsampling to 2000 cells maximum. RCTD was run in full mode (doublet_mode = “full”), enabling detection of multiple cell types per Visium spot (55μm diameter). RCTD performs platform-effect normalization to account for technical differences between scRNA-seq and spatial transcriptomics, followed by non-negative least squares (NNLS) regression to estimate cell type proportions. Normalized cell type weights (0-1, summing to 1 per spot) were extracted and visualized using scatterpie plots. Cell type proportions were compared across timepoints using Kruskal-Wallis tests with pairwise Wilcoxon post-hoc tests (Bonferroni correction, p < 0.05).

### Pathway activity scoring and spatial visualization

To quantify pathway activities at the spot level, we used UCell (v2.12.0) with pathway definitions harmonized to KEGG through msigdbr (v24.1.0; C2 CP:KEGG). Short-chain fatty acid (SCFA) metabolism (propionate and butyrate) and key metabolic pathways relevant to kidney physiology and injury were emphasized across figures. UCell scores were visualized using spatial feature plots that employed consistent color scaling, and, where applicable, standardized sample ordering to facilitate visual comparison across conditions and regions.

### Pseudobulk analyses, PCA, and mean squared error distances

To summarize expression at higher granularity, we computed pseudobulk profiles by averaging spot-level expression within biologically meaningful groups (for example, by cell type and timepoint or by sample). PCA was then performed on scaled pseudobulk matrices (19,973 genes x 8 samples after variance filtering) to visualize dominant axes of variation. The first two principal components (PC1 and PC2) accounted for 45.9% and 15.9% of total variance respectively, with PC1 primarily reflecting injury phase-dependent metabolic rewiring and PC2 capturing homeostatic states (top 20 contributing genes per component are listed in Supplementary table 2). Pseudobulk PCA figures are presented with square aspect ratios. To quantify expression similarity between pseudobulk profiles, we computed pairwise mean squared error (MSE) distances and visualized these distances as square heatmaps with publication-scale typography.

### Correlation analyses among pathway programs

We evaluated relationships among metabolic, repair, and cell-death programs by computing Spearman correlations between UCell pathway scores. Where required by specific figure panels, we constrained the axis content (for example, restricting one axis to cycling/repair signatures or including ketogenesis/ketone-body and apoptosis/necroptosis programs). Correlation matrices were visualized with Complex Heatmap (v2.24.0), using coherent annotations and titles to facilitate interpretation across related figures.

### Spatial cell–cell communication analysis

Cell–cell communication was inferred using CellChat (v2.2.0). We supplied spot-level expression matrices together with spatial coordinates derived from the Seurat object. Analyses were conducted on the whole-sample level and stratified by anatomical regions to investigate potential region-specific signaling patterns. Relative information flow was summarized across signaling pathways.

### Human dataset analyses

The atlas was imported into Seurat, and KEGG-harmonized pathway scores were computed using UCell. Pathway activities across disease categories and cell types were visualized using UMAPs and violin plots. For rendering efficiency of very large cohorts, we applied stratified subsampling that preserved disease composition, ensuring visual clarity without distorting biological representation.

### Cell culture

Primary PT cells were generated from the Immorto-mouse crossed with Tgfbr2^fl/fl^ mice as previously described^59^. Briefly, PT cells were isolated from male mice and grown at 33°C in DMEM/F12 supplemented with 2.5% fetal bovine serum, hydrocortisone, insulin, transferrin, selenium, triiodothyronine, penicillin/streptomycin (complete PT media) and with IFNγ (companies, locations). Prior to experiments, PT cells were moved to 37°C and IFNγ removed to induce differentiation. PT cells were plated at 30% confluency in complete PT media and treated 4 days with 1 mM propionate (Sigma) or 0.1 mM butyrate (Sigma).

### Analyses of cell metabolism

For XFe Seahorse experiments, Mitostress tests, cell density optimization and working concentration titters for each inhibitor were performed following the manufacturer’s guidelines (Agilent). For Mito Stress tests, cells were treated with 1 mM propionate or 0.1 mM butyrate, and 25,000 cells were plated per well of a Seahorse plate one day before the measurement. The injection ports included 2 µM oligomycin, 2.5 µM carbonyl cyanide 4-(trifluoromethoxy) phenylhydrazone (FCCP) and 0.5 µM rotenone/antimycin, respectively (Agilent). After each experiment, cells were lysed using 1X passive lysis buffer (Promega, location) and Bradford assays (company, location) were performed to determine protein concentrations. All experiments were analyzed using Wave software.

### mRNA quantification

RNA from PT cells was isolated using the Nucleospin RNA extraction kit following the manufacturer’s instructions (Macherey Nagel, location). cDNA was generated using AffinityScript Multi-Temp Reverse Transcription (RT) kit (Agilent). RT-qPCR was performed using a MX3000p real-time PCR cycler (Agilent). Relative mRNA expression was determined by the ΔΔCT method, using p0 as a reference gene. Primer sequences were as follows (forward and reverse) p0: 5’-CTCGCTTTCTGG AGGGTGTC-3’ and 5’-CCGCAAATGCAGATGGATCA-3’; Pck1: 5’-ATGAAAGGCCGCACCATGTA-3’ and 5’-CCTCCAGCACAGATATGCCC-3’; Gk: 5’-CTCTCATAGCCTGAA AGCTGG-3’ and 5’-CGTGTTTTTGGCCTGTCCAT-3’; Pc: 5’-GCAGGGCGGAGCTAACATC-3’ and 5’-GGCTTATACTCCAGACGCCG-3’; Hmgcs1: 5’-TGATCCCCTTTGGTGGCTGAA-3’ and 5’-AGCTGTGTGAAGGACAGAGAAC-3’; Hmgcs2: 5’-TCAGTGGAAGCAAGCTGGAAA-3’ and 5’-CATCAACCGAGCCAGGGATT-3’; Bh2: 5’-TGTGCCGCAGGATCCACTA-3’ and 5’-GGACTCGAGTTTGAATACCTCGG-3’.

### Immunoblotting

PT cells were lysed with 25 mM Tris-HCl pH 8.0, 1 mM EDTA, 150 mM NaCl, 1% NP-40, 1 mM Na_3_VO_4_ and freshly added protease inhibitor cocktail (1 mM PMSF, 1 µg/ml aprotinin, 1 µg/ml leupeptin and 1 µg/ml pepstatin A). Lysates were separated by SDS-PAGE and proteins electrotransferred to nitrocellulose membranes. All primary antibodies were dissolved in 5% BSA and incubated with membranes overnight at 4 °C. The following primary antibodies were used: total OXPHOS (ab110413; Abcam), β-actin (Sigma) and polyclonal goat anti-mouse-HRP (31430; Pierce). Following addition of HRP substrate (NEL113001EA; PerkinElmer, location), chemiluminescence was recorded with a LAS-400 mini–Luminescent Image Analyzer (FUJIFILM) and quantified using Image J.

### Statistical analysis

For group comparisons nonparametric tests (Wilcoxon or Kruskal–Wallis, as appropriate) were used, and for correlation analyses Spearman’s rho was applied. Where multiple hypothesis testing was applied, P values were adjusted using the Benjamini–Hochberg method. Analyses were performed in R v4.5.0 with Seurat v5.3.1.1000, UCell v2.12.0, ComplexHeatmap v2.24.0, CellChat v2.2.0, msigdbr v24.1.0, SeuratDisk v0.0.0.9021, zellkonverter v1.18.0, and qs2 v0.1.5. Figures were rendered with ggplot2-based extensions, ensuring fixed aspect ratios for embeddings. Unpaired Student’s *t* test was used to compare two sets of data, and one-way ANOVA followed by multiple comparisons was used to compare 3 or more groups, with p < 0.05 considered statistically significant. The number of independent experiments is indicated in each figure legend, and results are presented as mean ± SEM.

## Data availability

Spatial transcriptomics raw FASTQ files and metadata are available at GEO accession GSE310547 (https://www.ncbi.nlm.nih.gov/geo/). Processed data are available via an interactive website (https://fgcz-sushi.uzh.ch). Source data will be provided with this paper. All other relevant data supporting the key funding of this study are available within the article and its Supplementary information files.

## Code availability

All analyses were performed based on the Sushi uzh/sushi: SUSHI: Supporting User for SHell script Integration (github.com) and ezRun uzh/ezRun: An R meta-package for the analysis of Next Generation Sequencing data (github.com). Codes to reproduce all parts of the analysis can be requested at https://fgcz-sushi.uzh.ch

## Acknowledgments

This work was supported by the Swiss National Science Foundation (AMBIZIONE grant PZ00P3_179916 and the Swiss Federal Government Excellence Scholarship; NCCR “Kidney.CH” junior grant to S.N.K); the Swiss National Science Foundation (310030_184813 and NCCR “Kidney.ch”, 183774, to R.H.W); the French National Agency for Research (ANR-25-CPJ1-0134-01) to S.N.K S.N.K conceived the project and designed experiments. J.G, S.H, K.S and S.N.K performed the in vivo and in vitro experiments. S.N.K and P.G performed spatial transcriptomics data analysis. R.H.W supported the project and discussed the results. S.N.K supervised the project and wrote the manuscript with input from all authors.

## Competing Interests

None of the authors has conflicts of interest.

## Figure Legend

**Extended Data Figure 1**: Dot plots depicting cell type marker gene expression across robust cell type decomposition (RCTD) predicted cell types (a), proximal tubules (b), distal tubules (c), glomeruli (d), stromal/immune cells (e), and injured proximal tubules (f). DCT, distal convoluted tubule cells; Endo, endothelial cells; IC, collecting duct intercalated cells; Injured_PT1, type 1 injured proximal tubule cells; Injured_PT2, type 2 injured proximal tubule cells, LOH, loop of Henle cells; Macro, macrophages; Macro_prolif, proliferating macrophages; Neu, neutrophiles; T_prolif, proliferating T cells; PC, collecting duct principal cells; Pod, podocytes; PT, proximal tubule cells; S1/S2/S3, segments 1/2/3; Severe injured, severely injured PT; Stromal, stromal cells.

**Extended Data Figure 2:** Differential renal cell type proportion analyses across conditions. Box plots showing cortical cell type proportions across uninjured (n = 1), peak acute injury (3W, n = 2), transition (5W, n = 3) and chronic injury (8W, n = 2) phases. DCT, distal convoluted tubule cells; Endo, endothelial cells; IC, collecting duct intercalated cells; Injured_PT1, type 1 injured proximal tubule cells; Injured_PT2, type 2 injured proximal tubule cells, LOH, loop of Henle cells; Macro, macrophages; Macro_prolif, proliferating macrophages; Neu, neutrophiles; T_prolif, proliferating T cells; PC, collecting duct principal cells; Pod, podocytes; PT, proximal tubule cells; S1/S2/S3, segments 1/2/3; Severe injured, severely injured PT; Stromal, stromal cells.

**Extended Data Figure 3:** Decreased metabolic pathways and increased inflammatory pathways upon kidney injury compared to the uninjured state. (a-c) Dot plots showing significantly upregulated KEGG metabolic pathways in uninjured kidney compared to peak acute injury (a), transition (b) and chronic injury (c) phases. (e-f) Dot plots showing significantly downregulated KEGG metabolic pathways in uninjured kidney compared to peak acute injury (d), transition (e) and chronic injury (f) phases.

**Extended Data Figure 4**: UCell metabolic pathway enrichment analyses per condition and cell type. (a) UCell score heatmap showing main substrates and oxidative metabolic pathway enrichment in uninjured and injured (3W, 5W and 8W) kidneys. (b) UCell score heatmap showing main substrates and oxidative metabolic pathways enrichment in cells of the PT, DCT, LOH and PC. DCT, distal convoluted tubule cells;; Injured_PT1, type 1 injured proximal tubule cells; Injured_PT2, type 2 injured proximal tubule cells, LOH, loop of Henle cells; PC, collecting duct principal cells; PT, proximal tubule cells; S1/S2/S3, segments 1/2/3.

**Extended Data Figure 5**: Impaired keto-gluconeogenic metabolic pathways post-ischemia reperfusion injury. UCell violin plots showing Peroxisome (a), Val Leu Ile degradation (b), Propanoate metabolism (c), Butanoate metabolism (d), Tryptophan metabolism (e), PPAR signaling (f) pathway enrichments in sham operated controls (CTRL) and 4h, 12h 2 days, 14 days and 6 weeks post-ischemic injury mouse kidneys from GSE139107 dataset. Comparisons were statistically assessed using Wilcoxon rank-sum tests (*, p < 0.05; **, p < 0.01; ***, p < 0.001).

**Extended Data Figure 6:** UCell enrichment of fatty acid metabolism and short chain fatty acid metabolism per cell type. Violin plots showing fatty acid metabolism (a) and propionate metabolism (b) enrichment per cell type across conditions (uninjured, 3W, 5W and 8W injury phases). DCT, distal convoluted tubule cells; Injured_PT1, type 1 injured proximal tubule cells; Injured_PT2, type 2 injured proximal tubule cells, LOH, loop of Henle cells; PC, collecting duct principal cells; PEC, parietal epithelial cells, PT, proximal tubule cells; S1/S2/S3, segments 1/2/3.

**Extended Data Figure 7:** Sub-clustering analysis of the LOH cell cluster. (a) UMAP showing 4 clusters (g0, g1, g2 and g3) generated through Harmony integration. (b) Heatmap showing cluster specific marker expression. (c) UMAPs showing expression of TAL markers within the four identified clusters. (d) UMAPs showing metabolic pathway enrichment within the 4 clusters. (e) Violin plots showing metabolic pathway enrichment within each cluster.

**Extended Data Figure 8**: Propionate and butyrate promote the expression of renal ketogenic and gluconeogenic enzymes. (a) Table showing top 10 enriched metabolic pathways based on 463 genes characterizing the transition phase. (b) Clustergram showing genes involved in enriched metabolic terms in the transition phase. (c) RNA profiling generated by the mouse ENCODE project, showing the renal specificity of some ketogenic (Bdh2 and Oxct1) and gluconeogenic (glycerol kinase) enzymes involved in enriched metabolic pathways based on 463 genes characterizing the transition phase. RPKM, reads per kilobase million. (d) mRNA quantification by RT-qPCR showing the effect of 1 mM propionate (Pr) and 0.1 mM butyrate (Bu) on gluconeogenic enzymes (Pck1, Gk and Pc), and on ketogenic enzymes (Hmgcs1, Hmgcs2 and Bdh2) in PT cells.

**Extended Data Figure 9:** M1 and M2 macrophage analyses. UCell score Violin plots showing M1 and M2 enrichment in Macro_prolif and other_Macro (upper panel) and Macro_prolif and other_Macro distribution based on M1/M2 signature (lower panel). Macro, macrophages; Macro_prolif, proliferating macrophages.

**Extended Data Figure 10:** Keto-gluconeogenic metabolic pathway changes per cell type. UCell violin plots showing Peroxisome (a), propanoate metabolism (b), butanoate metabolism (c), tryptophan metabolism (d) and valine, leucine and isoleucine degradation pathway enrichments per cell type in healthy controls (normal), AKF and CKD kidney biopsies from integrated single-cell atlas of healthy and injured human kidney biopsies, enabling comparisons across disease states (GSE121862). Endo, endothelial cells; PT, proximal tubule cells; Fib, fibroblasts; CNT, connecting tubule cells, DCT, distal convoluted tubule cells; TAL, thick ascending limb cells; tDL, thin descending limb cells; Peri, pericytes; PC, collecting duct principal cells; IC, collecting duct intercalated cells. Comparisons were statistically assessed using Wilcoxon rank-sum tests (*, p < 0.05; **, p < 0.01; ***, p < 0.001).

## Notes

https://www.ncbi.nlm.nih.gov/geo/query/acc.cgi?acc=GSE310547

https://fgcz-sushi.uzh.ch

### Competing Interest Statement

The authors have declared no competing interest.

