## Extended Data Figure 1 for "Keto-gluconeogenic metabolic axis mirrors renal homeostasis and post-injury response in spatial transcriptomics"

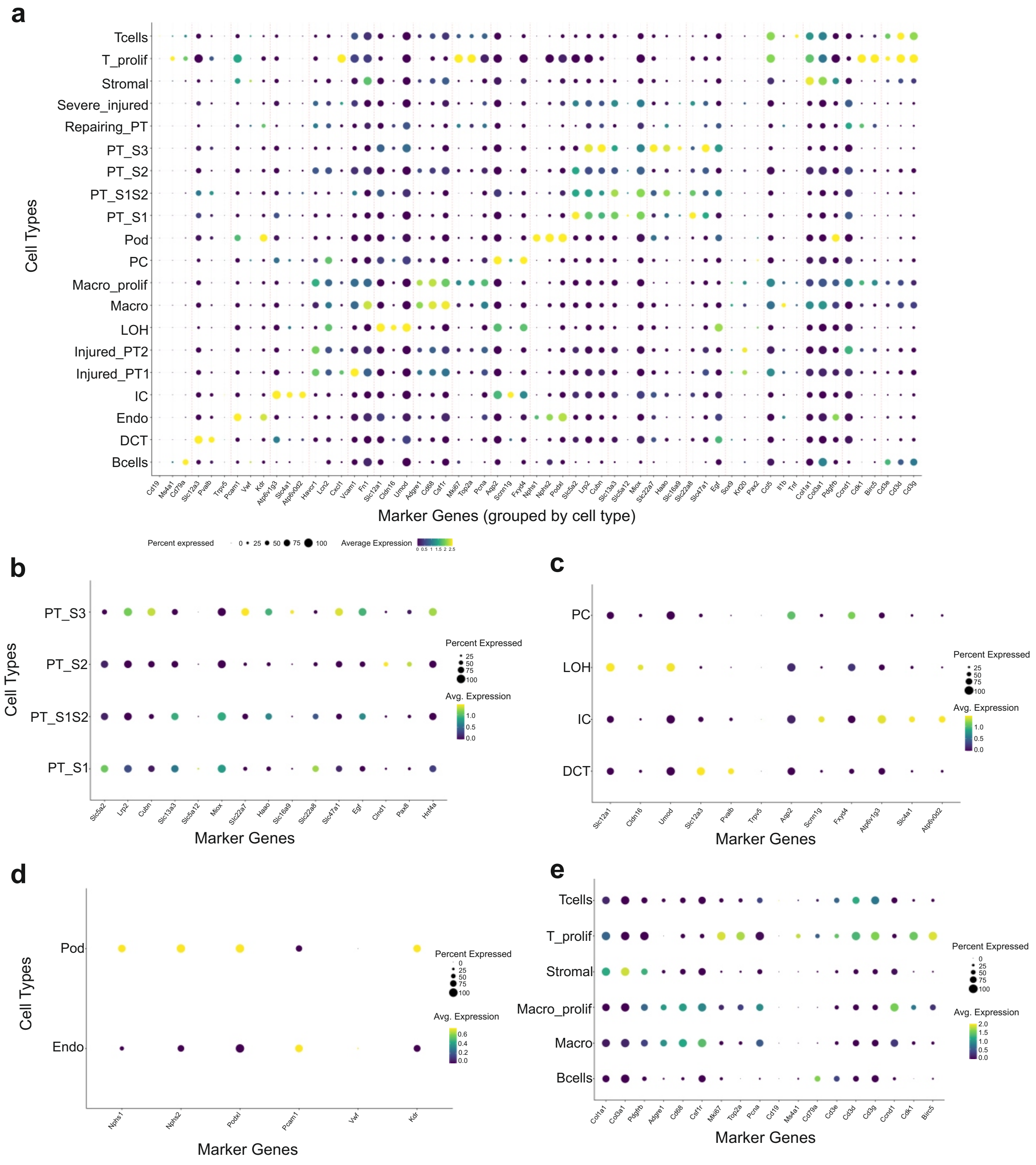


**Extended Data Figure 1**: Dot plots depicting cell type marker gene expression across robust cell type decomposition (RCTD) predicted cell types (a), proximal tubules (b), distal tubules (c), glomeruli (d), stromal/immune cells (e), and injured proximal tubules (f). DCT, distal convoluted tubule cells; Endo, endothelial cells; IC, collecting duct intercalated cells; Injured_PT1, type 1 injured proximal tubule cells; Injured_PT2, type 2 injured proximal tubule cells, LOH, loop of Henle cells; Macro, macrophages; Macro_prolif, proliferating macrophages; Neu, neutrophiles; T_prolif, proliferating T cells; PC, collecting duct principal cells; Pod, podocytes; PT, proximal tubule cells; S1/S2/S3, segments 1/2/3; Severe injured, severely injured PT; Stromal, stromal cells.
