## Extended Data Figure 2 for "Keto-gluconeogenic metabolic axis mirrors renal homeostasis and post-injury response in spatial transcriptomics"

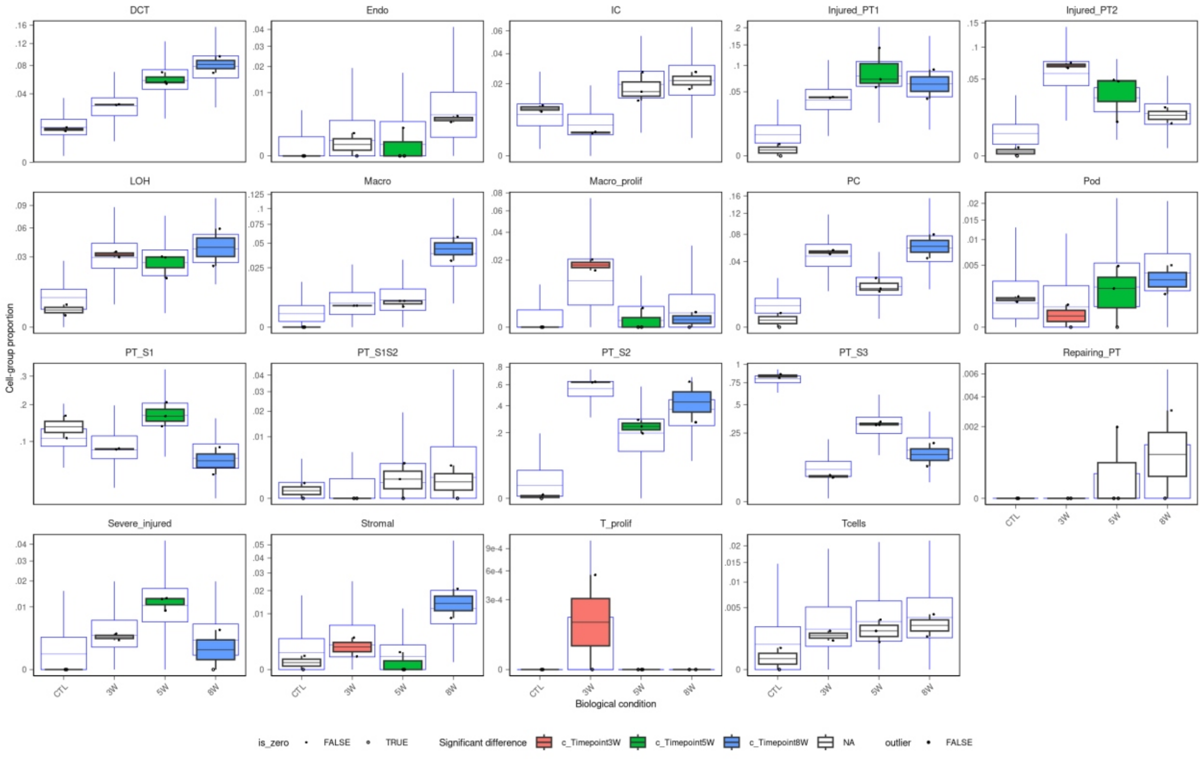


**Extended Data Figure 2:** Differential renal cell type proportion analyses across conditions. Box plots showing cortical cell type proportions across uninjured (n = 1), peak acute injury (3W, n = 2), transition (5W, n = 3) and chronic injury (8W, n = 2) phases. DCT, distal convoluted tubule cells; Endo, endothelial cells; IC, collecting duct intercalated cells; Injured_PT1, type 1 injured proximal tubule cells; Injured_PT2, type 2 injured proximal tubule cells, LOH, loop of Henle cells; Macro, macrophages; Macro_prolif, proliferating macrophages; Neu, neutrophiles; T_prolif, proliferating T cells; PC, collecting duct principal cells; Pod, podocytes; PT, proximal tubule cells; S1/S2/S3, segments 1/2/3; Severe injured, severely injured PT; Stromal, stromal cells.
