## Extended Data Figure 3 for "Keto-gluconeogenic metabolic axis mirrors renal homeostasis and post-injury response in spatial transcriptomics"

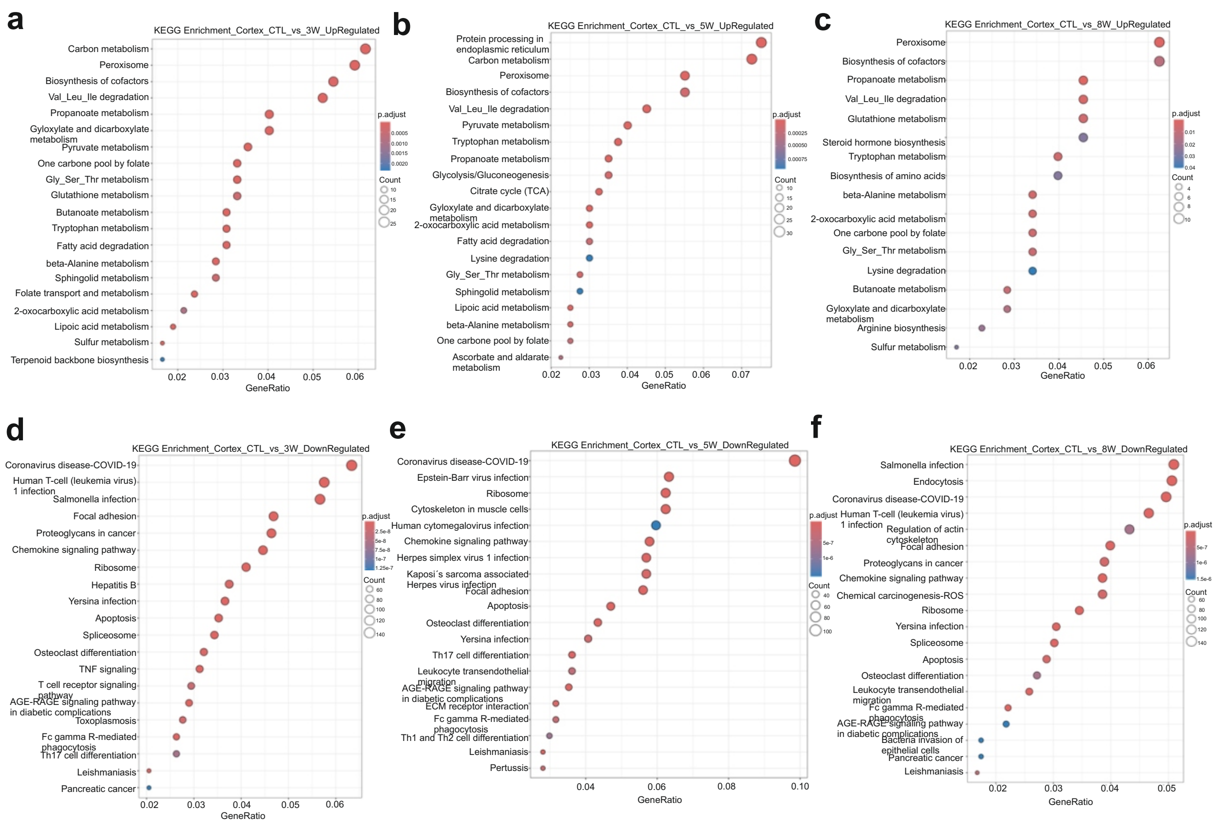


**Extended Data Figure 3:** Decreased metabolic pathways and increased inflammatory pathways upon kidney injury compared to the uninjured state. (a-c) Dot plots showing significantly upregulated KEGG metabolic pathways in uninjured kidney compared to peak acute injury (a), transition (b) and chronic injury (c) phases. (e-f) Dot plots showing significantly downregulated KEGG metabolic pathways in uninjured kidney compared to peak acute injury (d), transition (e) and chronic injury (f) phases.
