## Extended Data Figure 4 for "Keto-gluconeogenic metabolic axis mirrors renal homeostasis and post-injury response in spatial transcriptomics"

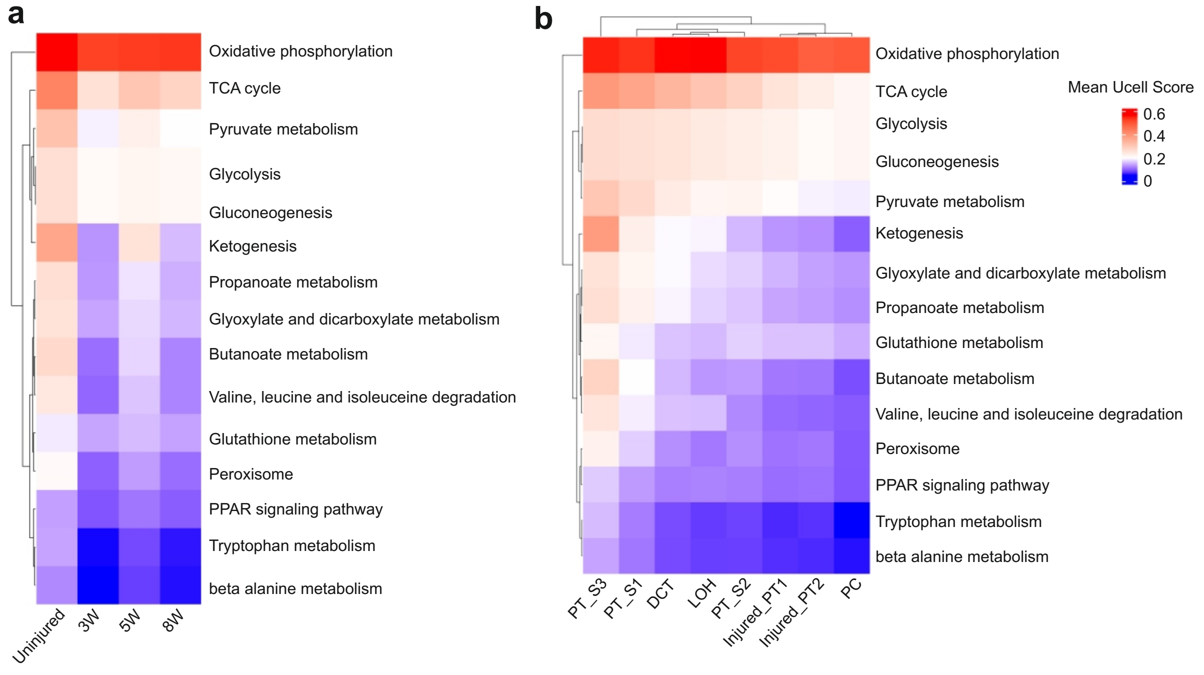


**Extended Data Figure 4:** UCell metabolic pathway enrichment analyses per condition and cell type. (a) UCell score heatmap showing main substrates and oxidative metabolic pathway enrichment in uninjured and injured (3W, 5W and 8W) kidneys. (b) UCell score heatmap showing main substrates and oxidative metabolic pathways enrichment in cells of the PT, DCT, LOH and PC. DCT, distal convoluted tubule cells;; Injured_PT1, type 1 injured proximal tubule cells; Injured_PT2, type 2 injured proximal tubule cells, LOH, loop of Henle cells; PC, collecting duct principal cells; PT, proximal tubule cells; S1/S2/S3, segments 1/2/3.
