## Extended Data Figure 5 for "Keto-gluconeogenic metabolic axis mirrors renal homeostasis and post-injury response in spatial transcriptomics"

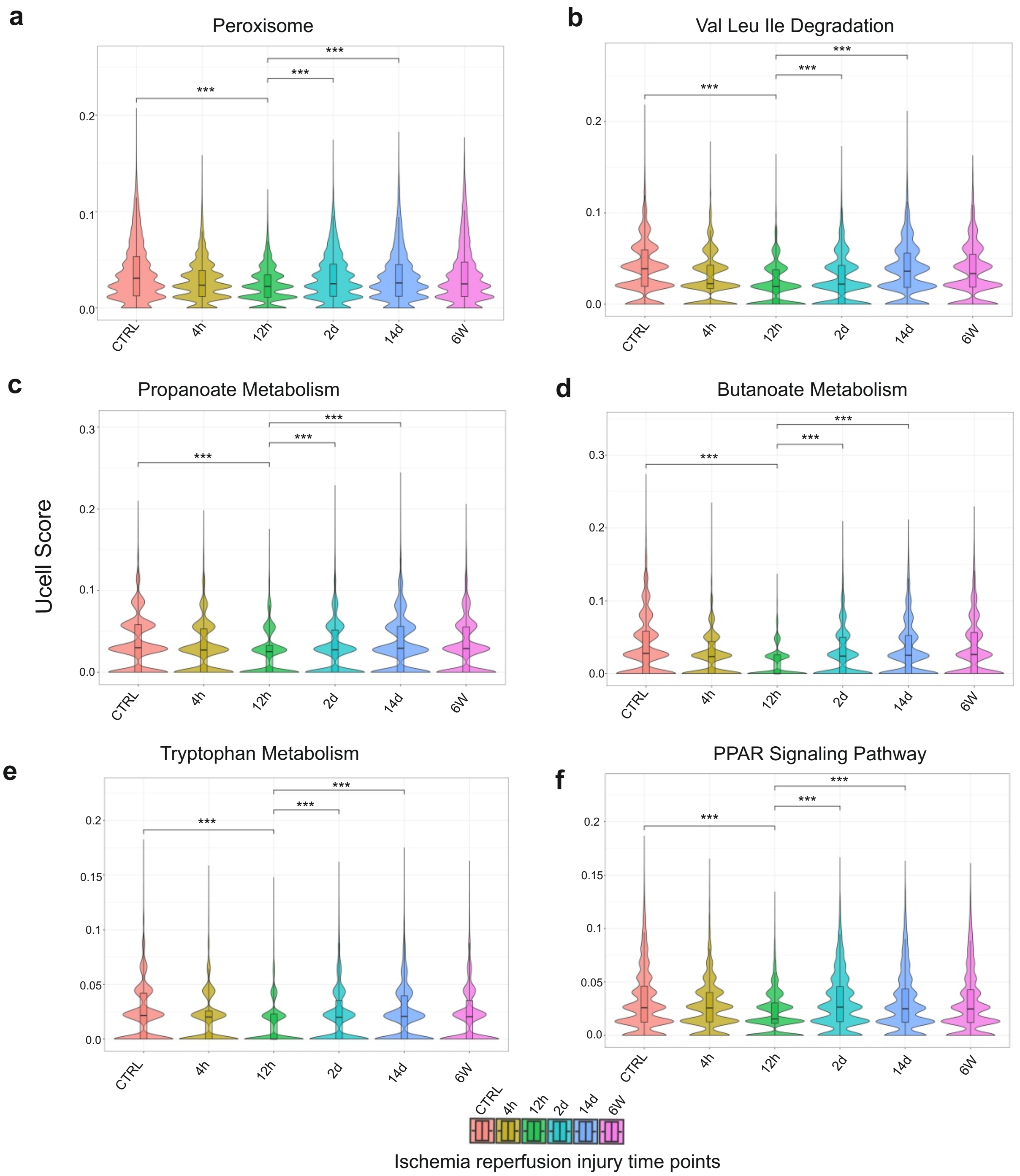


**Extended Data Figure 5**: Impaired keto-gluconeogenic metabolic pathways post-ischemia reperfusion injury. UCell violin plots showing Peroxisome (a), Val Leu Ile degradation (b), Propanoate metabolism (c), Butanoate metabolism (d), Tryptophan metabolism (e), PPAR signaling (f) pathway enrichments in sham operated controls (CTRL) and 4h, 12h 2 days, 14 days and 6 weeks post-ischemic injury mouse kidneys from GSE139107 dataset. Comparisons were statistically assessed using Wilcoxon rank-sum tests (*, $p < 0.05$; **, $p < 0.01$; ***, $p < 0.001$).
