## Extended Data Figure 6 for "Keto-gluconeogenic metabolic axis mirrors renal homeostasis and post-injury response in spatial transcriptomics"

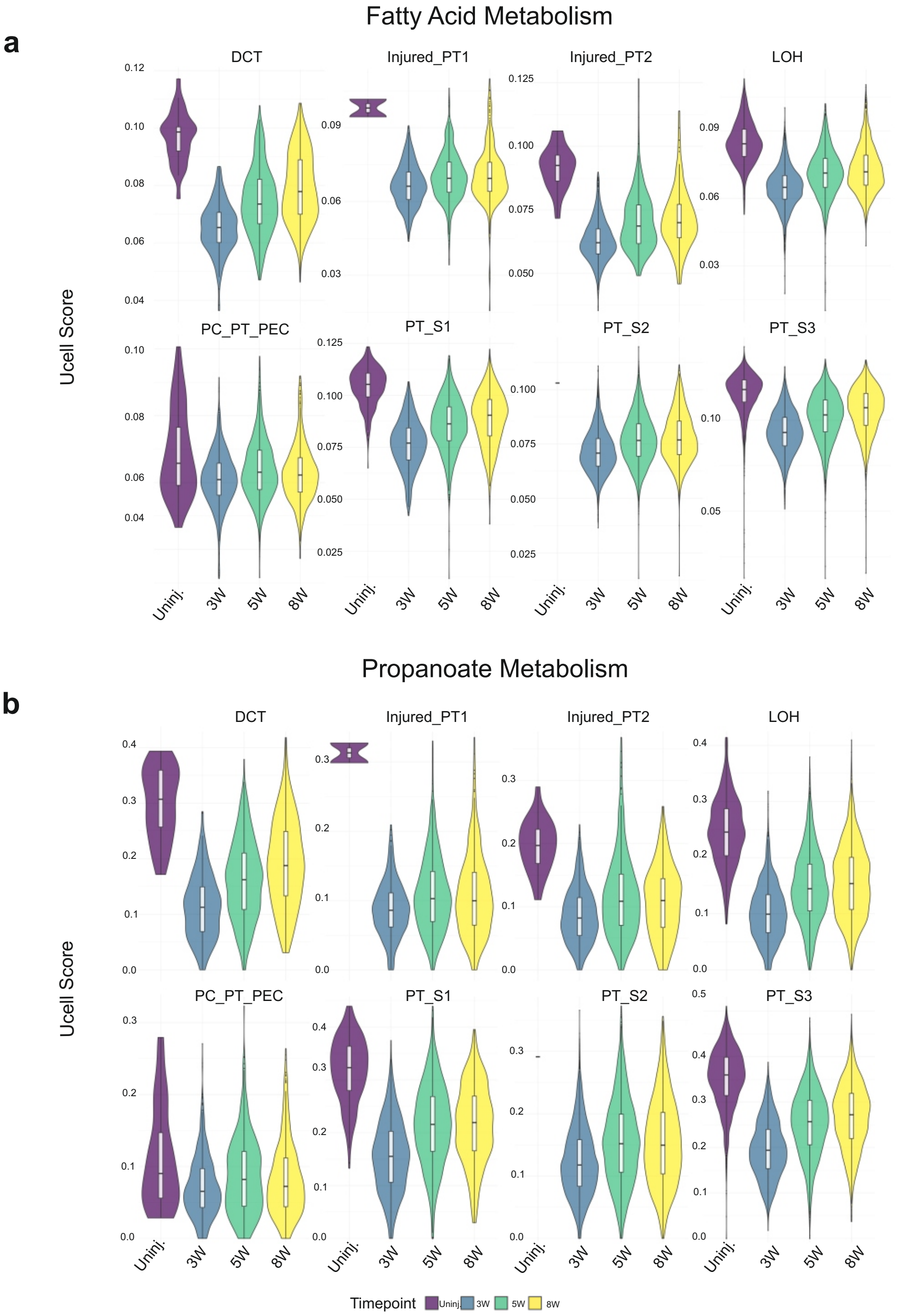


**Extended Data Figure 5**: UCell enrichment of fatty acid metabolism and short chain fatty acid metabolism per cell type. Violin plots showing fatty acid metabolism (a) and propionate metabolism (b) enrichment per cell type across conditions (uninjured, 3W, 5W and 8W injury phases). DCT, distal convoluted tubule cells; Injured_PT1, type 1 injured proximal tubule cells; Injured_PT2, type 2 injured proximal tubule cells, LOH, loop of Henle cells; PC, collecting duct principal cells; PEC, parietal epithelial cells, PT, proximal tubule cells; S1/S2/S3, segments 1/2/3.
