## Extended Data Figure 7 for "Keto-gluconeogenic metabolic axis mirrors renal homeostasis and post-injury response in spatial transcriptomics"

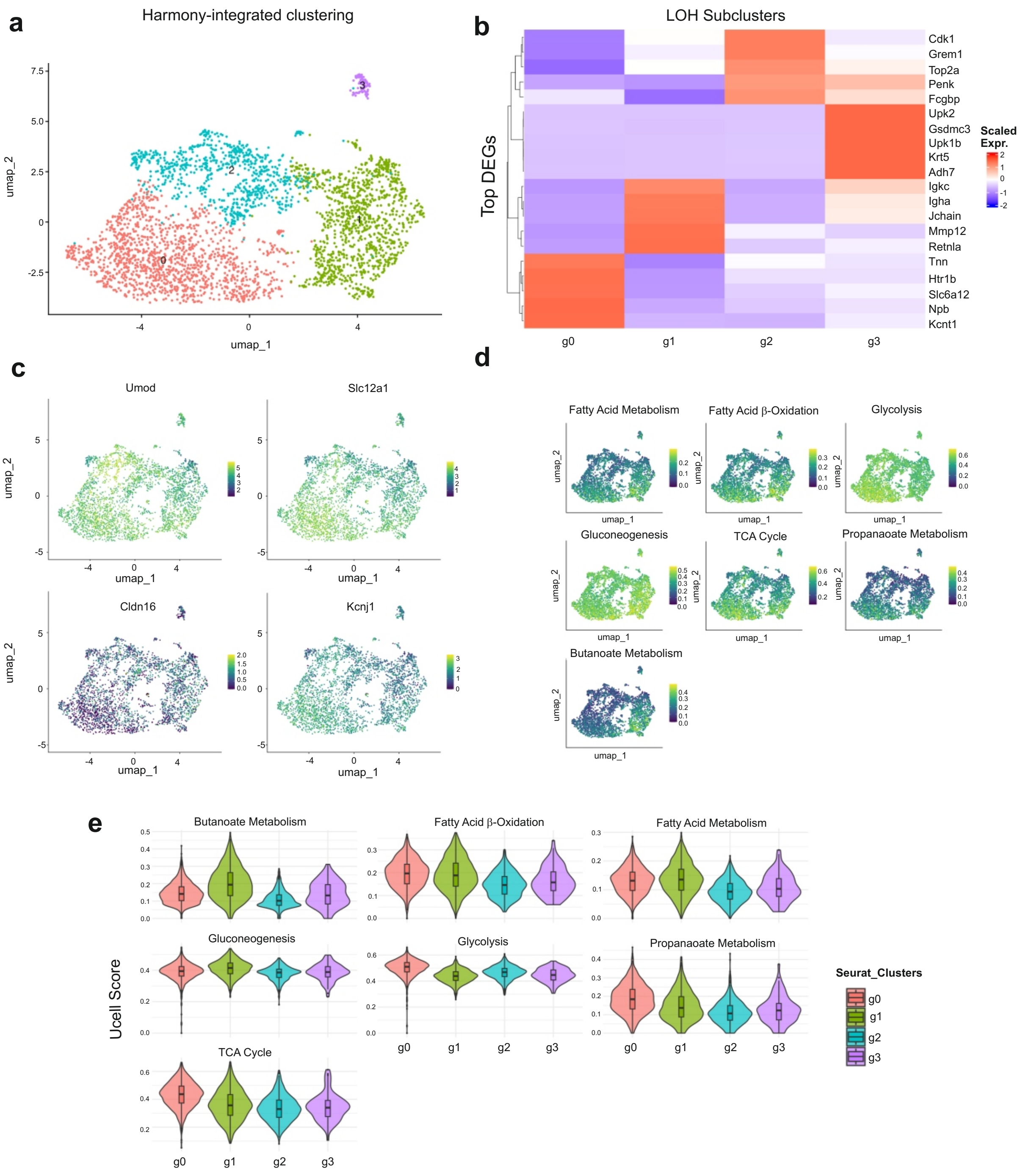


**Extended Data Figure 6:** Sub-clustering analysis of the LOH cell cluster. (a) UMAP showing 4 clusters (g0, g1, g2 and g3) generated through Harmony integration. (b) Heatmap showing cluster specific marker expression. (c) UMAPs showing expression of TAL markers within the four identified clusters. (d) UMAPs showing metabolic pathway enrichment within the 4 clusters. (e) Violin plots showing metabolic pathway enrichment within each cluster.
