## Extended Data Figure 8 for "Keto-gluconeogenic metabolic axis mirrors renal homeostasis and post-injury response in spatial transcriptomics"

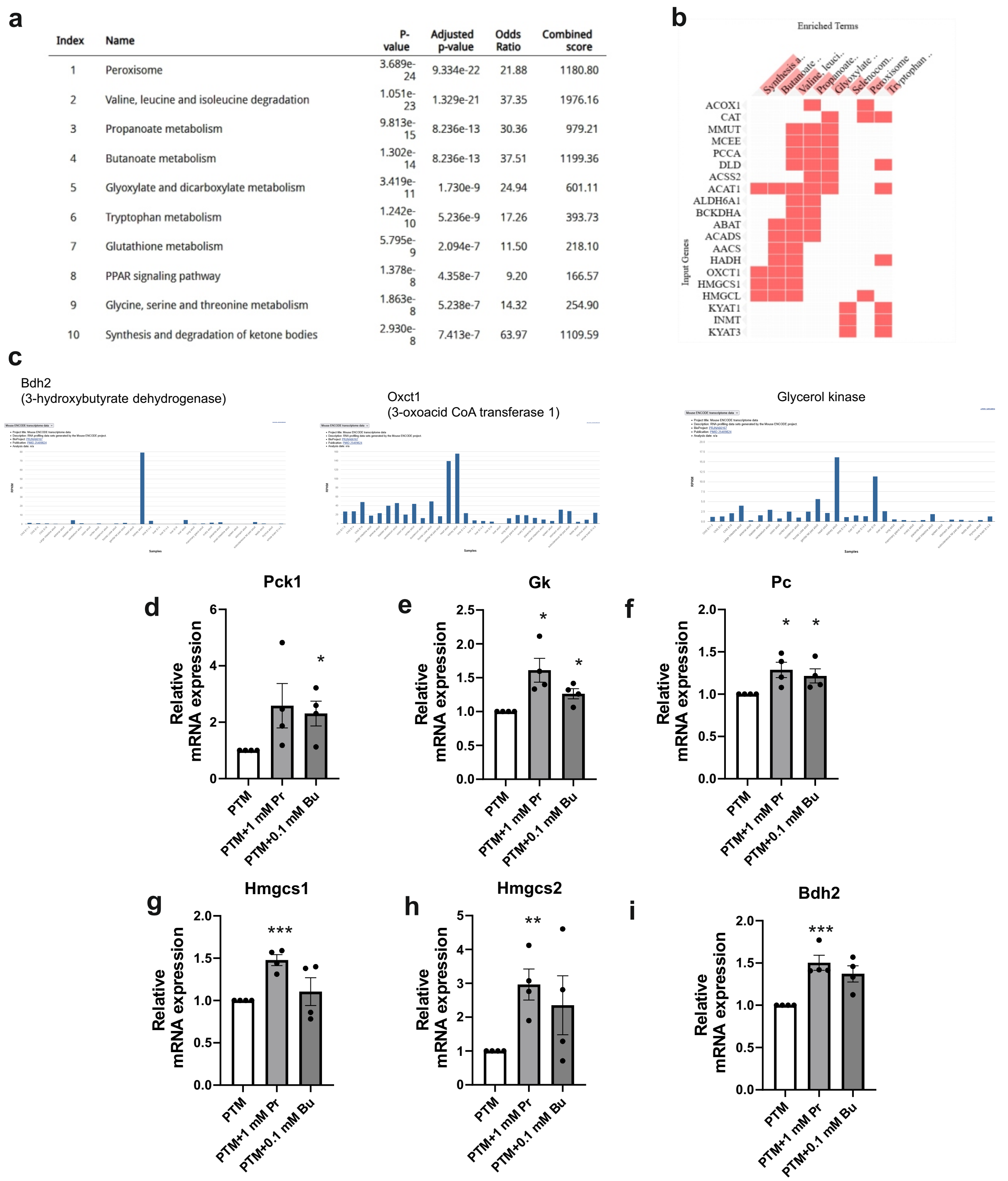

| Index | Name | P-value | Adjusted p-value | Odds Ratio | Combined score |
|---|---|---|---|---|---|
| 1 | Peroxisome | 3.689e-24 | 9.334e-22 | 21.88 | 1180.80 |
| 2 | Valine, leucine and isoleucine degradation | 1.051e-23 | 1.329e-21 | 37.35 | 1976.16 |
| 3 | Propanoate metabolism | 9.813e-15 | 8.236e-13 | 30.36 | 979.21 |
| 4 | Butanoate metabolism | 1.302e-14 | 8.236e-13 | 37.51 | 1199.36 |
| 5 | Glyoxylate and dicarboxylate metabolism | 3.419e-11 | 1.730e-9 | 24.94 | 601.11 |
| 6 | Tryptophan metabolism | 1.242e-10 | 5.236e-9 | 17.26 | 393.73 |
| 7 | Glutathione metabolism | 5.795e-9 | 2.094e-7 | 11.50 | 218.10 |
| 8 | PPAR signaling pathway | 1.378e-8 | 4.358e-7 | 9.20 | 166.57 |
| 9 | Glycine, serine and threonine metabolism | 1.863e-8 | 5.238e-7 | 14.32 | 254.90 |
| 10 | Synthesis and degradation of ketone bodies | 2.930e-8 | 7.413e-7 | 63.97 | 1109.59 |



**Extended Data Figure 7 :** Propionate and butyrate promote expression of renal enzymes ketogenic and gluconeogenic enzymes. (a) Table showing top 10 enriched metabolic pathways based on 463 gene input characterizing the transition phase, their p value, adjusted p-value, Odds ratio and combined score. (b) Clustergram showing some genes involved in enriched metabolic terms in the transition phase. (c) RNA profiling generated by the Mouse ENCODE project showing renal specificity of some ketogenic (Bdh2 and Oxct1) and gluconeogenic (Gycerol kinase) enzymes involved in enriched metabolic pathways based on 463 gene input characteristic of the transition phase. RPKM (Reads per kilobase million). (d, e, f) Preliminary qpcr analyses showing the effect of 1mM propionate (Pr) and 0.1mM of butyrate (Bu) on mRNA expression of gluconeogenic enzymes (Pck1, Gk and Pc). (g, h, i) Preminary qpcr analysis showing the effect of 1mM propionate (Pr) and 0.1mM of butyrate (Bu) on mRNA expression of ketogenic enzymes (Hmgcs1, Hmgcs2 and Bdh2).
