## Extended Data Figure 9 for "Keto-gluconeogenic metabolic axis mirrors renal homeostasis and post-injury response in spatial transcriptomics"

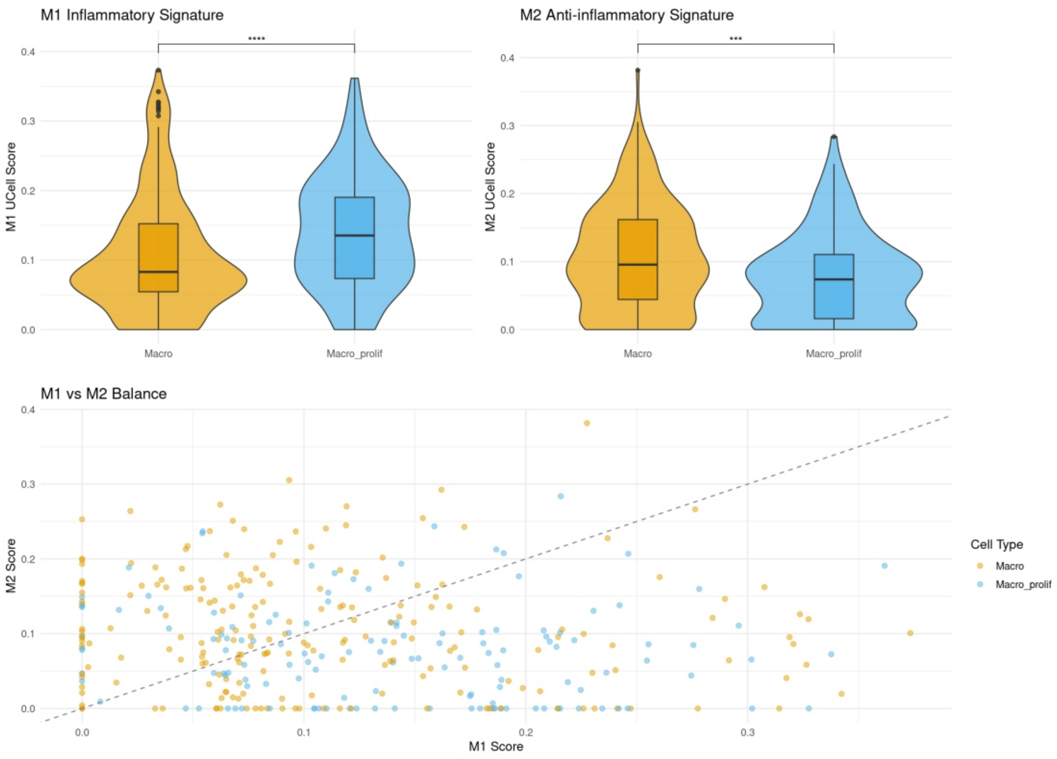


**Extended Data Figure 8 :** M1 and M2 macrophage analyses. UCell score Violin plots showing M1 and M2 enrichment in Macro_prolif and other_Macro (upper panel) and Macro_prolif and other_Macro distribution based on M1/M2 signature (lower panel). Macro, macrophages; Macro_prolif, proliferating macrophages.
