## Extended Data Figure 10 for "Keto-gluconeogenic metabolic axis mirrors renal homeostasis and post-injury response in spatial transcriptomics"

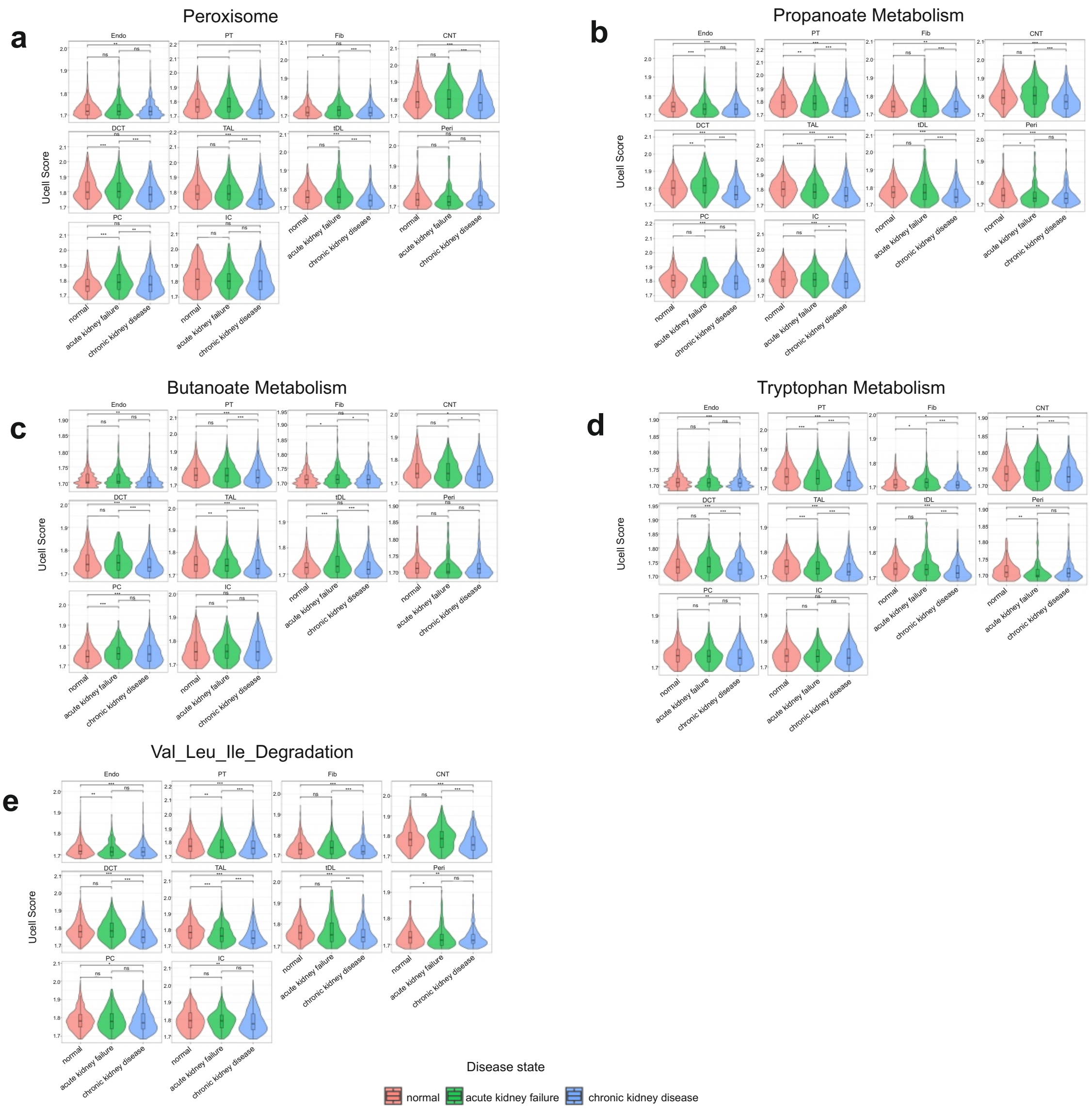


**Extended Data Figure 9:** Keto-gluconeogenic metabolic pathway changes per cell type. UCell violin plots showing Peroxisome (a), propanoate metabolism (b), butanoate metabolism (c), tryptophan metabolism (d) and valine, leucine and isoleucine degradation pathway enrichments per cell type in healthy controls (normal), AKF and CKD kidney biopsies from integrated single-cell atlas of healthy and injured human kidney biopsies, enabling comparisons across disease states (GSE121862). Endo, endothelial cells; PT, proximal tubule cells; Fib, fibroblasts; CNT, connecting tubule cells, DCT, distal convoluted tubule cells; TAL, thick ascending limb cells; tDL, thin descending limb cells; Peri, pericytes; PC, collecting duct principal cells; IC, collecting duct intercalated cells. Comparisons were statistically assessed using Wilcoxon rank-sum tests (*, $p < 0.05$; **, $p < 0.01$; ***, $p < 0.001$).
